# A single BRCA2 BRC repeat supports viability while multiple repeats ensure resilience under stress

**DOI:** 10.64898/2026.01.07.698272

**Authors:** Arun Prakash Mishra, Sounak Sahu, Satheesh Sengodan, Gemma Moore, Swati Priya, Natasha Oberoi, Julia R. Jensen, Eileen Southon, Mellissa Galloux, Dylan Caylor, Teresa Sullivan, Sandra S. Burkett, Mary E. Albaugh, Parirokh P. Awasthi, Francesco Tomassoni Ardori, Lino Tessarollo, Ryan B. Jensen, Shyam K. Sharan

## Abstract

BRC repeats are integral to BRCA2 function and mediate RAD51 loading during homologous recombination (HR). Their number varies across species, but mammals, including mice and humans, harbor eight repeats. Prior studies have suggested functional redundancy among these repeats, but the biological significance of maintaining multiple repeats remains unresolved. Here, we demonstrate that the presence of a single BRC repeat, either BRC2 or BRC4, is sufficient for mouse embryonic stem cell (mESC) viability, RAD51 loading, PARP inhibitor resistance and protection of stalled replication forks. Consistent with these findings, we show that knock-in mice with a single BRC repeat 2 or 4 are viable and exhibit normal growth and fertility. In contrast, embryonic fibroblasts from these mice display genomic instability and impaired RAD51 recruitment. Notably, this defect is rescued under low oxygen culture conditions, which mimics physoxic levels, whereas exposure to oxidative stress impairs RAD51 recruitment in mESCs harboring a single BRC repeat. Together, our findings indicate that while a single BRC repeat is sufficient under physiological conditions, the evolutionary retention of multiple BRC repeats likely ensures robust genome stability under extreme oxidative stress.

## Introduction

BRCA2 is a well-known tumor suppressor functioning as a genome caretaker by recruiting RAD51 to DNA double-strand breaks (DSB) facilitating their repair by homologous recombination (HR)^1,2^. BRCA2 is essential for the survival of normal cells, including mouse embryonic stem cells^3^. One of the key functional domains of human BRCA2 are the eight BRC repeats, spanning residues 1002–2085 (Fig. 1a). Across metazoans, the BRC repeats have undergone significant evolutionary diversification in both number and sequence. The majority of mammalian BRCA2, including human and murine, contain 8 BRC repeats, each comprising of 35 non-identical (but conserved)^4^ amino acids connected by a non-conserved linker. In contrast, BRCA2 protein from *Trypanosoma* species harbors between 1 and 15 BRC-like repeats (Suppl. Fig. 1a). The eight mammalian BRC repeats are known to physically interact with RAD51 at differing affinities, with BRC1-4 exhibiting higher RAD51 affinity than BRC5-8^5^. Structural analysis elucidated two highly conserved hydrophobic regions (FxxA and LFDE) within BRC repeats responsible for RAD51 interaction^4,6^. However, mutational analysis of human BRC4 in *Ustilago maydis* revealed the FxxA motif is functionally more important^7^.

**Fig. 1:**
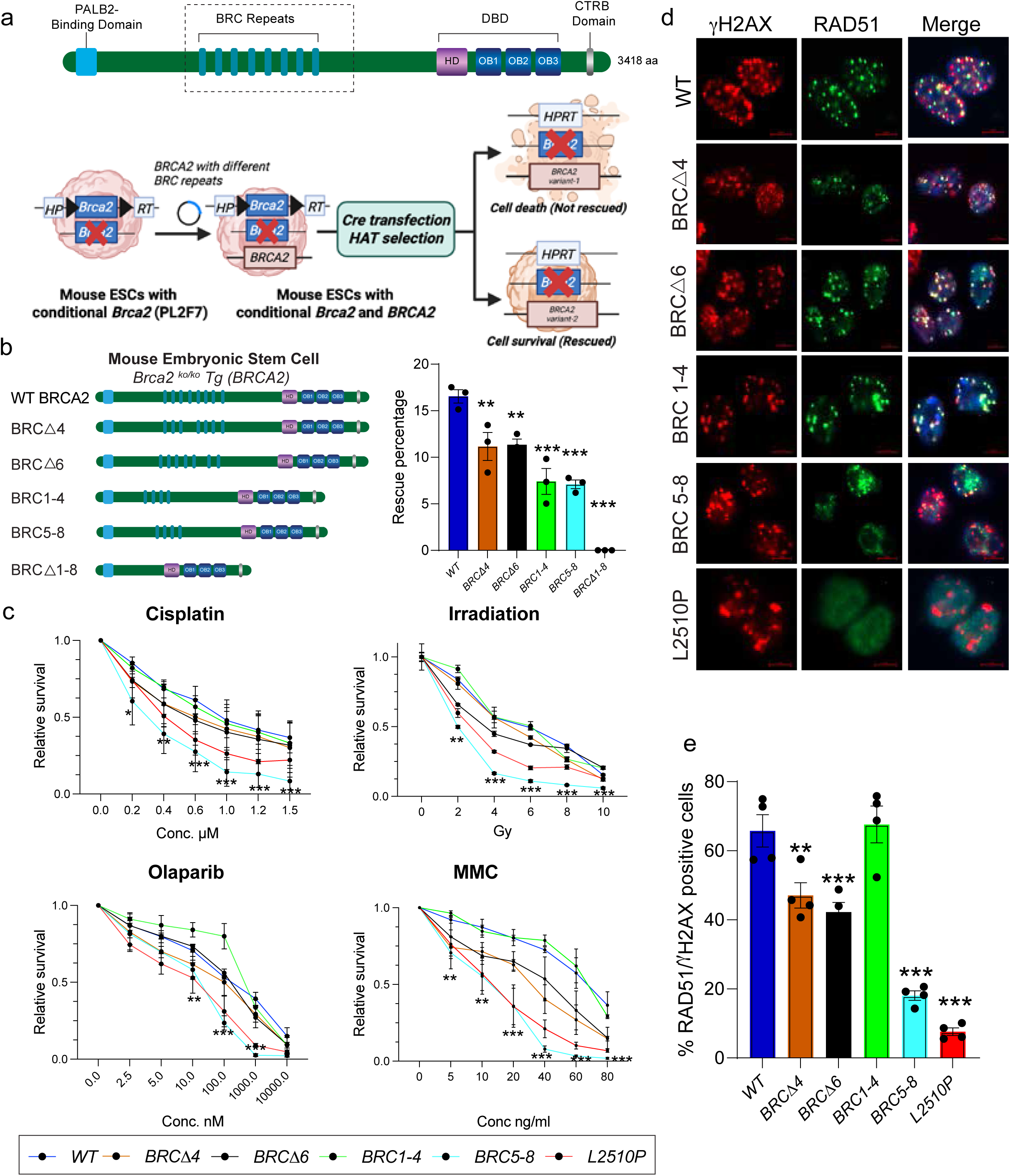
BRC repeats 5-8 are dispensable for BRCA2 functions. a) Pictorial representation of BRCA2 with different functional domains including N-terminal PALB2 binding domain, eight BRC repeat, C-terminal DNA binding domain (DBD) and the C-terminus RAD51 binding (CTRB) domain. Schematic representation of mESC-based BRCA2 functional complementation assay for cell survival using HAT selection. b) HAT rescue percentage of mESCs harboring BRCA2 with different BRC repeat deletions. BRCA2, with all the BRC repeats deleted, failed to rescue any mESC colony (n=3 independent clones, error bar-SEM, one way ANOVA, compared with WT). c) XTT based drug sensitivity assay using different DNA damaging agents. mESCs harboring BRCA2 with BRC5-8 are hypersensitive to all the drugs comparable to the known hypomorphic BRCA2 variant L2510P (n=3 independent clones, error bar-SEM, Students t-test, two tailed, for each point compared with WT). d) Representative images depicting RAD51 foci at the DSBs generated by 6Gy IR to mESCs harboring BRCA2 with different BRC repeat deletions. e) Quantification of percentage RAD51 positive nuclei observed in (d). mESCs with BRC5-8 exhibited significantly lower number of RAD51 positive nuclei compared to WT (n=4 independent clones, error bar-SEM, one way ANOVA, compared with WT). **p<0.01, ***p<0.001

Germline inheritance of a pathogenic variant in *BRCA2* significantly increases the lifetime risk of developing breast and ovarian cancers^8–10^. To date, most pathogenic missense variants have been identified in exon3, encoding the N-terminal Partner and Localizer of BRCA2 (PALB2) binding domain, and exons 15-26, encoding the C-terminal DNA binding domain (DBD) of BRCA2 (Fig.1a, top)^11,12^. While several frameshift and non-sense variants associated with increased cancer risk have been identified in exon11, that encodes the eight BRC repeats, no pathogenic missense variants have been reported in this exon^13^. A recent functional study has identified two hypomorphic variants, p.Ser1221Pro in BRC2 and p.Thr1980Iso in BRC7, that disrupt BRCA2’s interaction with RAD51 and render cells sensitive to chemotherapeutic agents^14^. The clinical significance of these variants, however, remains to be validated. Moreover, characterization of cisplatin and Poly(ADP-ribose) polymerase inhibitor (PARPi) resistant *BRCA2-*mutant breast (HCC1428) and pancreatic cancer cell line (Capan-1), along with patient tumors, has revealed revertant variants in *BRCA2* that restores, in *cis,* the open reading frame encoding two BRC repeats^15^. These studies hint towards inherent functional redundancy within BRC repeats. Moreover, a mini-BRCA2 carrying a single BRC repeat (repeats 1, 2, 3 or 4) exhibits HR proficiency, albeit two to five times lower than BRCA2 with BRC repeats 1-4 further corroborating their functional redundancy^16^.

Considering the inherent functional redundancy within BRC repeats, why species harbor multiple repeats is unknown. In this study we used mouse embryonic stem cells (mESC) and mouse models to investigate the relevance of presence of multiple BRC repeats. We demonstrate that the presence of a single BRC repeat is sufficient to support mESC viability and perform majority of canonical BRCA2 functions. Moreover, we were able to generate knock-in mice carrying BRCA2 with a single BRC repeat, BRC2 or BRC4. Both homozygous and hemizygous mutant mice are born at expected Mendelian ratios and are fully viable and fertile. However, contrary to the results observed in mESCs, mutant primary fibroblasts exhibited reduced RAD51 recruitment at DSBs. This RAD51 recruitment defect was alleviated under low oxygen (physoxic) conditions, whereas subjecting mESCs, harboring a single BRC repeat, to oxidative stress, by H_2_O_2_ treatment, impaired RAD51 recruitment. Our results demonstrate that oxidative stress destabilizes RAD51 interaction when BRCA2 contains a single BRC repeat, whereas the presence of multiple repeats confers functional robustness. These findings suggest a mechanistic basis by which multiple BRC repeats contribute to safeguarding the genome under oxidative stress.

## Results

### Functional evaluation of BRCA2 BRC repeat mutants in mouse ES cells

We evaluated the functional significance of BRCA2 BRC repeats using a well-established mESC-based functional assay^17^. We used PL2F7 mESC line in which one of the endogenous *Brca2* alleles is non-functional (knockout or *KO*) and the other is a conditional allele (*CKO*) flanked with two halves of human *HPRT* minigene (Fig. 1a)^17^. We electroporated PL2F7 mESCs with recombineered *BRCA2* cloned in a BAC (Bacterial artificial chromosome) having different BRC deletions, and deleted the endogenous conditional *Brca2* allele, to examine their impact on cell viability^23^. We confirmed loss of the conditional allele in viable HAT resistant clones by Southern analysis (Suppl. Fig. 1b-d).

BRC repeats 1-4 and 5-8 are known to be functionally distinct in RAD51 and ssDNA binding based on biochemical studies^18^. Therefore, we generated BAC clones containing only BRCA2 with BRC repeats 1-4 or 5-8 (BRC5-8 and BRC1-4 deleted, respectively) to examine their functional relevance. We also generated BRCA2 with deletion of the well-studied BRC4 (BRCΔ4) and the least conserved BRC repeat BRC6 (BRCΔ6). We obtained HAT resistant *Brca2-*null colonies, confirmed by Southern blot analysis (Suppl. Fig. 1c), that were rescued by BAC expressing full length (WT) *BRCA2* as well as by all other BRC mutants albeit with a reduced rescue rate (Fig. 1b). Interestingly, loss of all eight BRC repeats (BRCΔ1–8) failed to result in any viable clone (Fig. 1b).

Further, we assessed the sensitivity of mESCs expressing BRC mutant BRCA2 towards DNA-damaging agents by an XTT-based cell proliferation assay. mESCs expressing BRCA2 BRC5-8 were hyper-sensitive to all the DNA-damaging agents examined similar to the cells expressing a known pathogenic BRCA2 variant (p.Leu2510Pro, Fig. 1c). However, the sensitivities of cells expressing BRCA2 with BRC1-4, BRCΔ4 and BRCΔ6 were comparable to that of WT BRCA2.

Next, we examined the RAD51 recruitment ability of cells expressing BRCA2 with varying number of BRC repeats at the DSBs. We used γ−irradiation (IR) to induce DSBs and found mESCs expressing WT BRCA2 as well as BRCΔ4, BRCΔ6 and BRC1-4 mutants to have a comparable number of RAD51 foci (marker for HR) colocalizing with γH2AX (marker for DSBs) (Fig. 1d,e). However, the cells expressing BRCA2 with BRC5-8 exhibited a significantly reduced number of RAD51 foci that also appeared to be more diffused (Fig. 1d,e). As previously reported, BRCA2 L2510P mutant mESCs showed minimal or no IR-induced RAD51 foci^19^. These results indicated that cells expressing BRCA2 with BRC1-4 are proficient in BRCA2’s canonical functions (cell survival, resistance to DNA-damaging drugs and RAD51 recruitment) but those expressing BRCA2 with BRC5-8, although proficient in cell survival, lack the ability of RAD51 recruitment thereby rendering the cells sensitive to genotoxins.

### A single BRC is proficient in BRCA2 canonical functions

Since the cells expressing BRCA2 with BRC1-4 exhibited proficiency in BRCA2 functions, we next sought to determine which of these four BRC repeats is critical. We generated mESCs expressing BRCA2 with BRC1-3, BRC1-2 as well as those with a single BRC repeat 1, 2, 3 or 4. We found BRCA2 with a single BRC repeat rescued the mESC lethality due to *Brca2* loss (Suppl. Fig. 1d) but there was around 66% reduction in the rescue rate compared to WT BRCA2 (Fig. 2a). These findings suggested that BRCA2 containing even a single BRC repeat can support mESC viability, albeit with reduced efficiency.

**Fig. 2:**
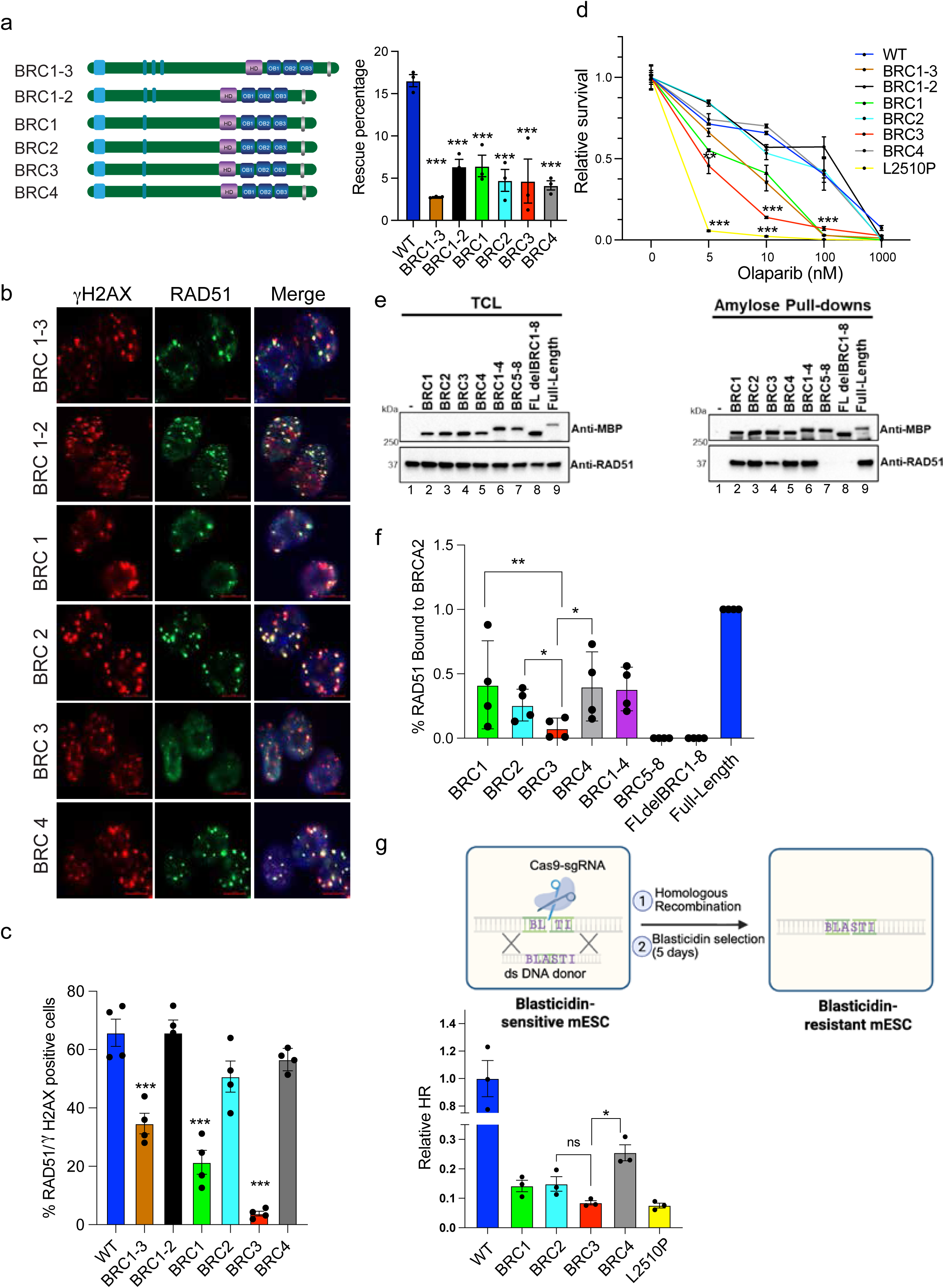
BRCA2 with single BRC repeat is proficient in canonical functions. a) HAT rescue percentage of mESCs with deletions in BRC repeat 1-4 (pictorial representation of mutant BRCA2 on left). Rescue percentage of BRCA2, with single BRC repeats, fell by one-third as compared to full length (n=3 independent clones, error bar-SEM, one way ANOVA, compared with WT). b) Representative images depicting RAD51 foci at the DSBs generated by 6Gy IR to mESCs harboring BRCA2 with single BRC repeat. c) Quantification of percentage RAD51 positive nuclei observed in (b). mESCs with BRC3 exhibited significantly lower number of RAD51 positive nuclei compared to WT and BRC2 and BRC4 were equivalent to that in WT (n=4 independent clones, error bar-SEM, one way ANOVA, compared with WT). d) Clonogenic survival assay to determine olaparib sensitivity of mESCs with deletions in BRC repeat 1-4. mESCs with BRC3 exhibited significant sensitivity towards olaparib (n=3 independent clones, error bar-SEM, Students t-test, two tailed). e) Western blots of total cellular lysates (TLC) from 293T cells transiently transfected with the indicated 2XMBP tagged BRCA2 BRC constructs (left panel, lanes 2-9). Lane 1 is a non-transfected control to visualize non-specific binding of RAD51 to the amylose beads. Western blots of amylose pull-downs from the same lysates depicted in first one. 36 hours post-transfection, cell lysates were detected by western blotting against MBP (BRCA2 constructs) or endogenous RAD51 (37 kDa). f) Quantification of band densitometry performed on amylose pull-down gel. RAD51 protein bound to each BRCA2 BRC construct was normalized to the amount of 2XMBP-BRCA2 fusion protein bound and eluted from amylose beads in the StainFree image. The percentage of RAD51 bound to full-length BRCA2 protein was set to 100% in the analysis (n=3 technical replicates, error bar-SEM, Students paired t-test). g) Schematic representation of blasticidin-resistance based HR assay in mESCs with single BRC repeats (top). Bottom panel shows quantification of HR levels relative to WT. Significant reduction in total HR is observed in all the samples with single BRC repeat. mESCs with BRC3 exhibited HR values comparable to hypomorphic BRCA2 mutation (L2510P) (n=3 independent clones, error bar-SEM, Students t-test, two tailed). *p<0.05, p<0.01, ***p<0.001

Further, we examined RAD51 recruitment in these mESCs after IR and observed a significantly reduced number of RAD51 foci in cells expressing BRCA2 with BRC1-3, BRC3 and BRC1 (Fig. 2b,c) as compared to WT mESCs. Remarkably, cells with either BRC2 or BRC4 exhibited RAD51 foci comparable to those observed in WT BRCA2 expressing cells (Fig. 2b,c).

We next performed clonogenic survival assay in the presence of varying doses of olaparib (PARP inhibitor) to determine the sensitivity of mESCs expressing mutant BRCA2 with various BRC repeats. Consistent with the observed reduction in RAD51 recruitment, cells expressing BRCA2 with BRC1-3, BRC1 and BRC3 resulted in significantly fewer colonies with olaparib treatment (Fig. 2d). Likewise, no significant difference in olaparib sensitivity was observed in cells expressing BRCA2 with BRC1-2, BRC2 and BRC4 compared to WT BRCA2. These findings suggest that BRCA2, with a single BRC repeat, BRC2 or BRC4, is proficient in RAD51 recruitment at the DSBs and provides resistance to olaparib.

To directly examine the binding of BRC mutant BRCA2 with RAD51 we performed pull-down experiments. We overexpressed recombinant 2XMBP tagged BRCA2 (full-length or indicated BRC repeat mutant) in HEK-293T cells and pulled down the fusion protein using amylose beads to quantify the amount of bound endogenous RAD51. We did not detect any RAD51 bound to BRCA2 deleted for BRC1-8 (Fig 2e, lane 8) or BRC1-4 (Fig. 2e, lane 7) in agreement with prior results^18^. Notably, significant amount of bound RAD51 was detected by full-length BRCA2 and BRCA2 containing BRC1-4 (Fig. 2e). The amount of RAD51 pulled down by BRCA2 with single BRC repeats 1, 2, 3 or 4 was reduced by at least 50% compared to the that with full-length protein (Fig. 2e,f, lanes 2-6, 9) with BRC3 showing the least RAD51 binding.

We next assessed the level of HR proficiency in the mESCs expressing WT as well as single BRC repeats (1,2,3 or 4). We utilized a previously described HR assay in mESCs carrying a 29bp deletion in the blasticidin resistance gene^19^ and repaired it with a donor DNA using CRISPR/Cas9. We quantified the number of blasticidin resistant colonies to assess HR efficiency. Surprisingly, despite normal RAD51 foci formation and olaparib resistance observed in BRC2/4 expressing cells, they exhibited significantly reduced HR efficiency. The level of HR in mESC with BRC3 was most reduced and comparable to the known BRCA2 hypomorph L2510P (Fig. 2g).

We also assessed the role of different BRC mutants in protection of stalled replication forks (RF) by performing DNA fiber assay^20^. The significance of BRC repeats in protecting stalled RFs is unknown as the C-terminal RAD51 binding (CTRB) domain of BRCA2, encoded by exon27 (Fig 1a, top), is known to play a crucial role^21^. We induced replicative stress by hydroxyurea (HU) to stall RF and found that all BRC mutants of BRCA2 are proficient in protecting stalled RFs (IdU/CldU ratio>0.9) suggesting that BRC repeats had no impact on this function (Suppl. Fig. 1e). A BRCA2 hypomorphic variant (R2336H) used as control exhibited defect in RF protection (IdU/CldU ratio <0.5)^22^.

### The FYSA residue is critical for BRC2-Mediated BRCA2 Function

We employed our previously developed CRISPR-based Saturation Genome Editing (SGE) approach in mESCs^23^ where we engineered a *Brca2*-null mESC line by stably expressing a transgene encoding human BRCA2 with a single BRC2 repeat [*Brca2*^−/−^; tg(*BRCA2^BRC2^*)]^24^. We used this approach (Fig. 3a) to confirm the functional redundancy among BRC repeats by individually integrating each BRC repeat (BRC1–8) in place of BRC2. In line with our previous results, we found BRC1/2/4 supported cell viability along with BRC8, whereas BRC3/5/6/7 alone failed to do so (Fig. 3b).

**Fig. 3:**
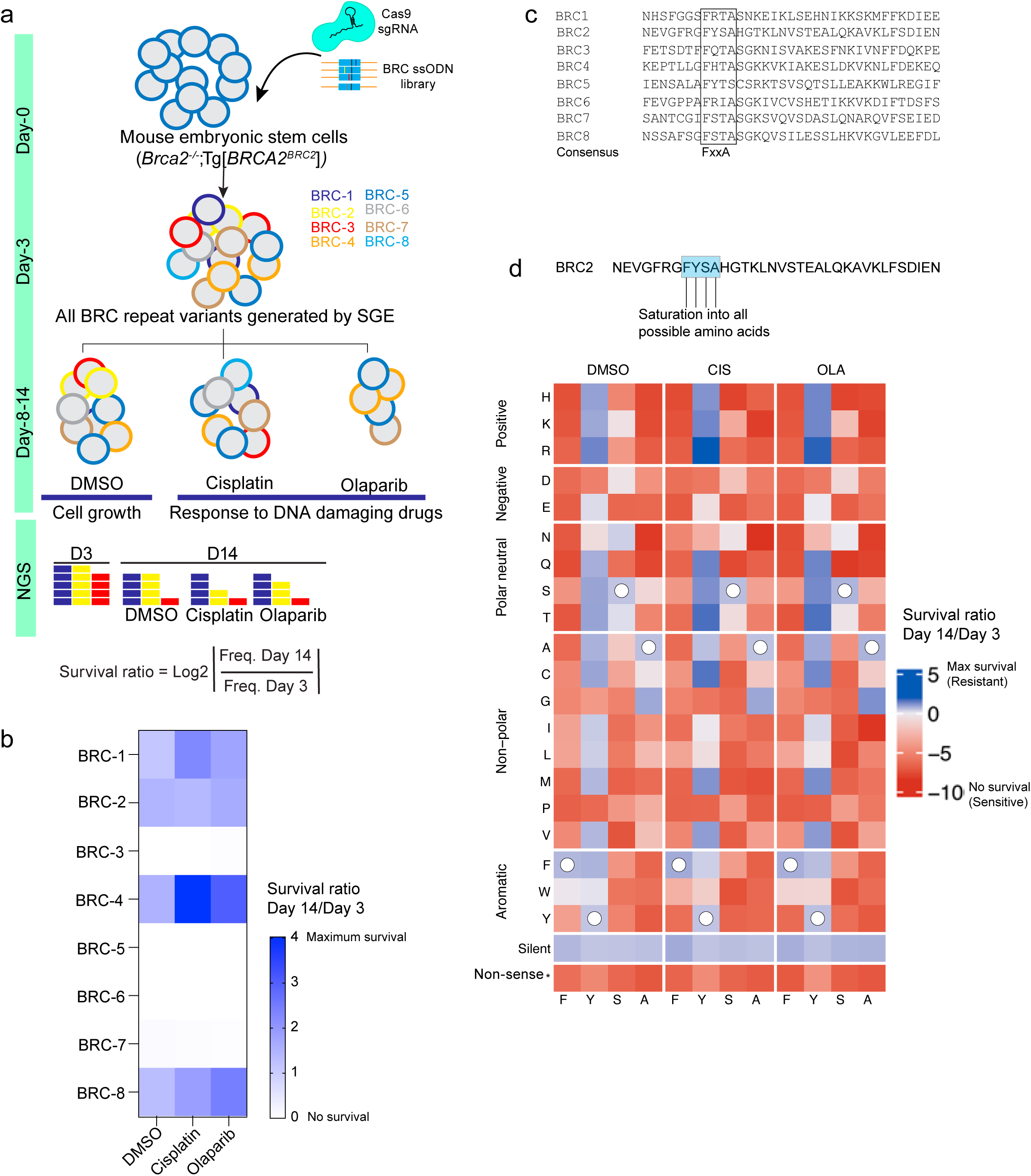
Functional analysis of BRC repeats in BRCA2. a) Schematic representation of the saturation mutagenesis approach using an ES cell line expressing BRCA2 only the BRC2 repeat. This approach was used to generate all 8 possible BRC repeat variants. WT BRC2 was targeted with a PAM modified BRC2 repeat. Cells were selected for 14 days, followed by deep sequencing to assess the impact of each BRC repeat on cell fitness and response to DNA-damaging agents. b) Heatmap showing cell survival in DMSO, cisplatin and olaparib for each of the eight BRC repeats. The shade of blue indicates relative survival, while white denotes lethality. Data is derived from three independent replicates. c) Sequence alignment showing the conserved FxxA motif across eight BRC repeats present in BRCA2. d) Heatmap showing saturation mutagenesis of the FYSA motif within the BRC2 repeat to all 20 amino acids. Data is derived from three independent replicates.

Since, BRC repeats harbor a conserved FxxA motif, which is essential for RAD51 binding (Fig. 3c)^4^, we next interrogated its functional importance by substituting the FYSA residues of BRC2 with all possible 20 amino acids at each position. We performed SGE in *Brca2*^−/−^; tg*BRCA2^BRC2^* mESC line (Fig. 3a). Analysis revealed that substitutions at F, S and A residues were largely intolerable, indicating their critical role in BRC2 function. Interestingly, conservative substitutions of F to W, I, L, Y, or V were partially tolerated in the absence of any drug treatment (Fig. 3d), consistent with findings in *Ustilago maydis*^7^. However, in the presence of cisplatin or olaparib, only W exhibited partial cell viability. In contrast, all substitutions of the tyrosine residue of FYSA, had only a moderate impact on cell viability. Our results reiterate that within the FxxA motif of a BRC repeat, F and A residues are critical for BRCA2 function.

### Generation of knock-in mice expressing BRCA2 with a single BRC repeat

Our studies in mESCs provide strong evidence that BRCA2 mutants with a single BRC repeat, especially BRC2 or BRC4, are fully functional. Encouraged by these findings, we tested whether BRCA2 with a single BRC repeat can support viability in mice, as loss of BRCA2 is embryonic lethal^3^. We targeted exon11 of mouse *Brca2*, which encodes eight BRC repeats, using CRISPR/Cas9 system with two different gRNAs, upstream and downstream of the eight BRC repeats (Suppl. Fig. 2a). Using two different donor oligos to replace BRC repeats 1-8 with either BRC2 or BRC4 alone, we obtained multiple correctly targeted knock-in mice (Suppl. Fig. 2b, c; Fig. 4a). We obtained heterozygous mice for the targeted alleles and will be referred to as *Brca2^BRC2/+^* (*BRC2/+,* for simplicity) and *Brca2^BRC4/+^* (*BRC4/+*). We intercrossed *BRC2/+* as well as *BRC4/+* mice to test whether mice expressing a single BRC repeat are viable in homozygous state. Remarkably, we obtained viable homozygous mice of both genotypes (*BRC2/BRC2* and *BRC4/BRC4*) in expected Mendelian ratios (Table 1). We also crossed the *BRC2/+* and *BRC4/+* mice with mice heterozygous for the *Brca2*-null allele (*Brca2^KO/+^*or *KO/+* for simplicity) to examine the impact of having a single BRC repeat in hemizygous state^3^. Notably, we also obtained hemizygous animals (*BRC2/KO* and *BRC4/KO*) in expected Mendelian ratios (Table1).

**Fig. 4:**
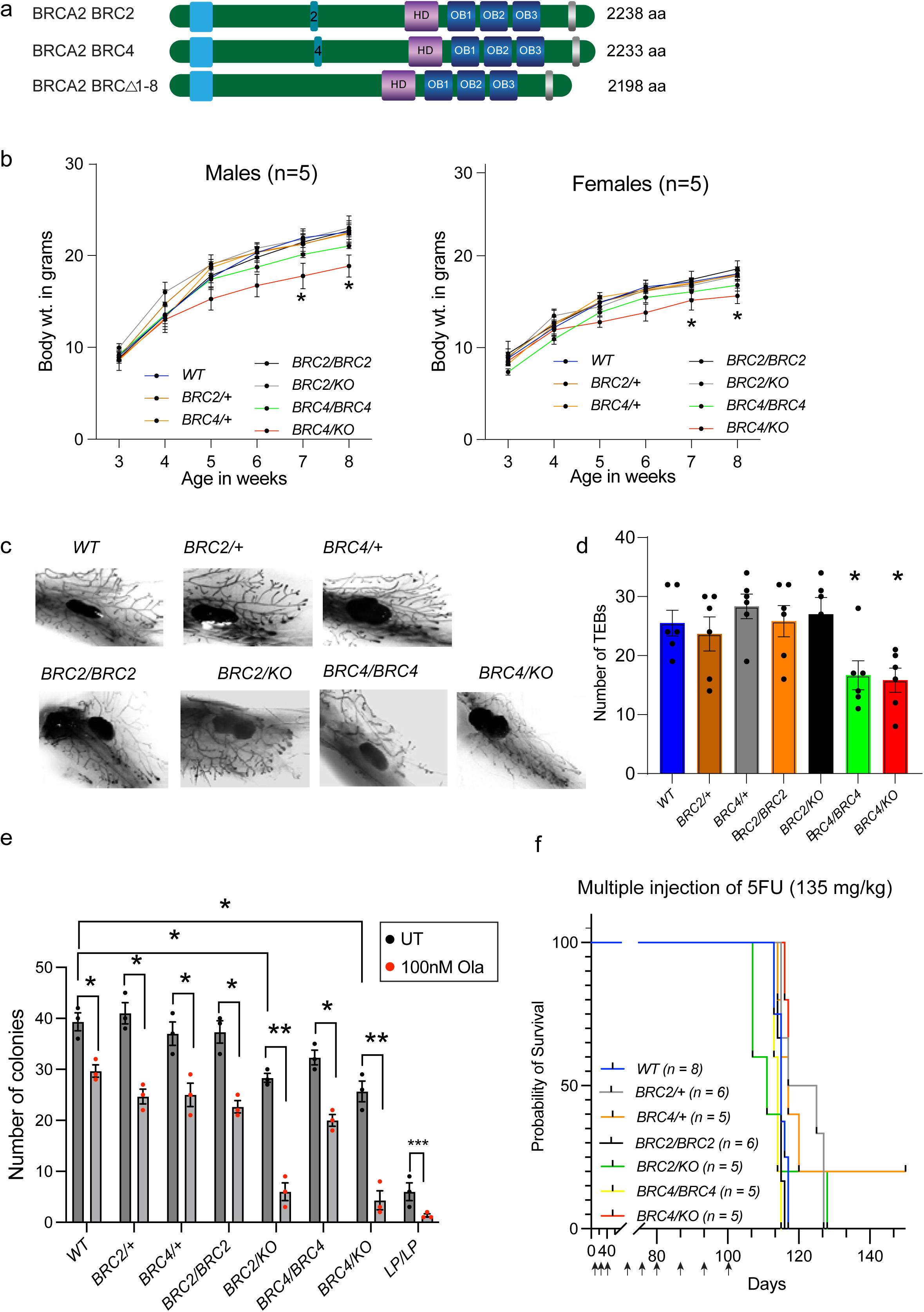
Phenotypic analysis of knock-in mice expressing BRCA2 with a single repeat, BRC2 and BRC4. a) Pictorial representation of BRCA2 protein expressing in the knock-in mice BRC2, BRC4 and BRCΔ1-8. b) Body weights of males and females of all the genotypes from weaning (3-weeks) to 8-weeks old. *BRC4/KO* mice showed lower body weights from week 5 onwards (n=5, error bars-SEM, Students t-test for each point compared with WT). c) Representative images of Carmine-alum-stained mammary glands, isolated from 5-week-old females of each genotype. d) Quantification of visible TEBs in the mammary glands. BRC4 homozygous and hemizygous females have significantly less TEBS compared to WT (n=6 mammary glands, error bar-SEM, one way ANOVA, compared with WT). e) Quantification of fetal liver colonies. *BRC2/KO* and *BRC4/KO* exhibited significantly lower number of colonies compared to WT. BRCA2 hypomorphic variant L2431P (*LP/LP*) is used as a negative control. *BRC2/KO* and *BRC4/KO* exhibited hypersensitivity to olaparib (n=3, error bar-SEM, one way ANOVA, compared with WT). f) Mouse survival plot after multiple 5FU injections at indicated time points to accelerate hematopoietic aging. No significant difference in survival is observed in any genotype (sample size mentioned in the parenthesis). *p<0.05, p<0.01, ***p<0.001

**Table 1.**
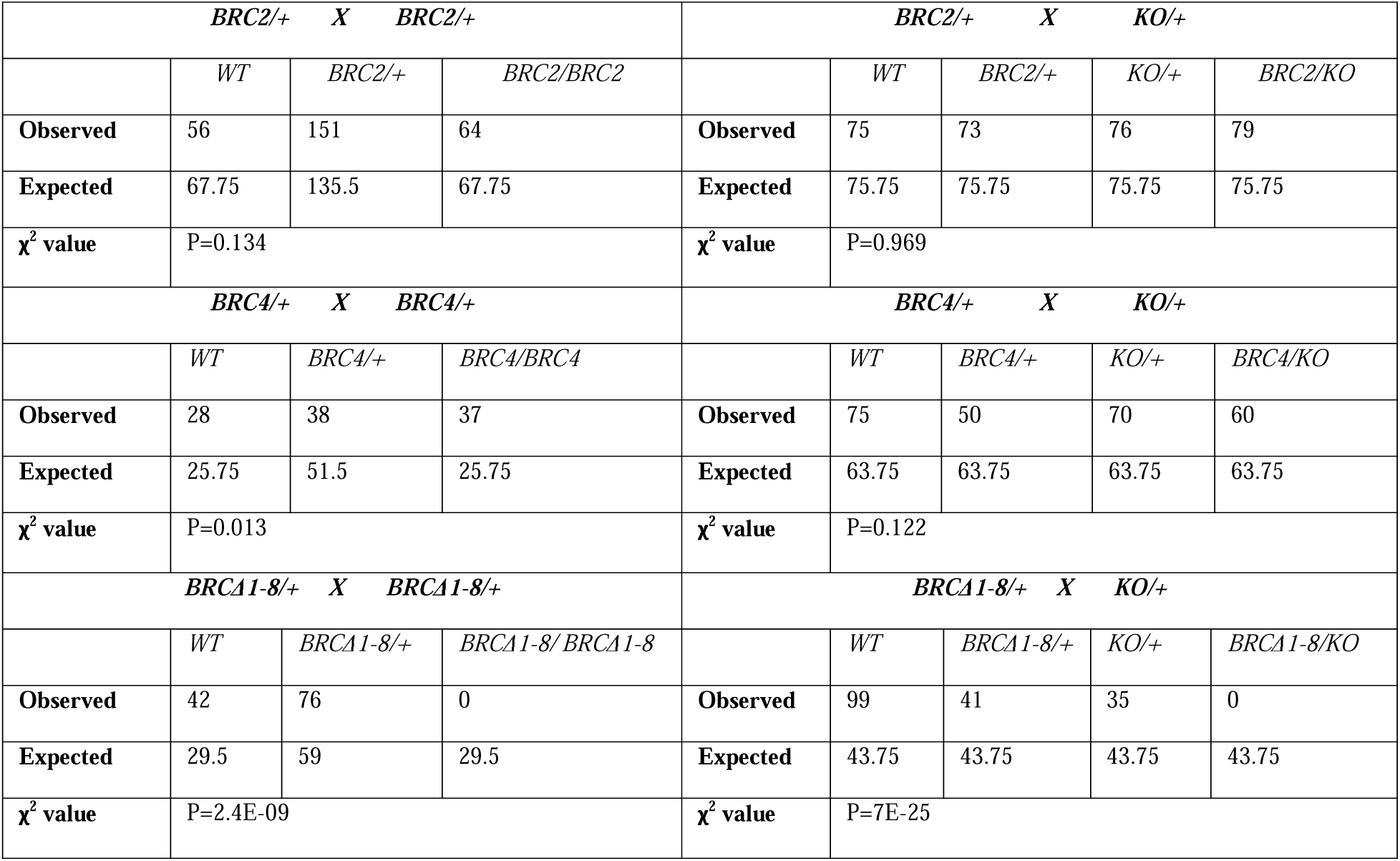
***BRC2* and *BRC4* mutant mice are obtained in expected Mendelian ratios in homozygous and hemizygous conditions**

In addition to the *BRC2* and *BRC4* knock-in alleles, we obtained a mouse that had an in-frame deletion of 3393bp (Suppl. Fig. 2c, Fig. 4a) in exon11 predicted to skip 1131aa of BRCA2 lacking all eight BRC repeats (*Brca2^BRC^*^Δ*1-8*^, referred to as *BRC*Δ*1-8*). When we intercrossed *BRC*Δ*1-8/+* mice or crossed them with *KO/+* mice, we failed to obtain any viable *BRC*Δ*1-8/BRC*Δ*1-8* or *BRC*Δ*1-8/KO* mice suggesting that loss of all the BRC repeats is embryonic lethal and at least one BRC repeat is required for embryonic development (Table 1). These results corroborate our *in vitro* findings showing that at least one BRC repeat of BRCA2 required to sustain viability.

### Mice harboring single BRC repeat of BRCA2 exhibit moderate developmental defects

We observed no overt phenotype in *BRC2* and *BRC4* homozygous as well as hemizygous mutant mice. However, starting five weeks postpartum, the *BRC4/BRC4* and *BRC4/KO* mice consistently displayed slightly reduced body weight compared with control mice, while the *BRC2/BRC2* and *BRC2/KO* mice maintained their body weight (Fig. 4b). To evaluate this growth defect, we examined mammary gland branching and quantified the number of terminal end buds (TEBs) as a proxy of developmental progression. We harvested mammary glands from 5-week-old females of various genotypes, stained them with carmine alum. While the number of TEBs in *BRC2/BRC2* and *BRC2/KO* females were comparable to the control genotypes, they were significantly reduced in *BRC4/BRC4* and *BRC4/KO* genotype (Fig. 4c, d).

The reduction in mammary TEBs could reflect a proliferation defect in mammary adult stem cells, and potentially other adult stem cells. Therefore, we examined fetal liver cells, the site of embryonic hematopoiesis, to assess defects in hematopoietic stem and progenitor cells (HSPCs). We isolated fetal liver cells from 16.5dpc embryos of each genotype and cultured them with growth factors to promote colony formation. Equivalent number of colony forming units (CFUs) were observed in control genotypes along with *BRC2* and *BRC4* homozygous mutants while *BRC2/KO* and *BRC4/KO* fetal liver cells exhibited mildly reduced number of CFUs (Fig. 4e, Suppl. Fig. 3a). Additionally, the hemizygous HSPCs were hypersensitive to olaparib treatment comparable to *LP/LP* genotype (a known HR-deficient p.Leu2431Pro BRCA2 variant expressing mouse)^19^ (Fig. 4e, Suppl. Fig. 3a). To examine the regenerative ability of HSPCs *in vivo*, we injected multiple (weekly) sub-lethal doses of 5-fluorouracil (5FU), to eradicate all dividing hematopoietic cells and stimulate the proliferation and differentiation of HSPCs^25^, in mice of all genotypes. As expected, repeated 5FU administration promoted exhaustion of proliferating HSPCs and mice become moribund after 100 days. However, all mice displayed a comparable pattern of mortality, and the mutants did not exhibit increased sensitivity to this proliferative stress (Fig. 4f). This result shows that despite a mild impairment in CFU formation in the *in vitro* assay, the hemizygous hematopoietic cells are functionally normal *in vivo*.

The minor developmental defects observed in BRC4 mutants did not affect their fertility and fecundity. Both male and female mutant mice were fully fertile, and we observed normal morphology in their testes and ovaries based on H&E staining (Suppl. Fig. 3b). Furthermore, spermatocyte spreads revealed normal meiotic progression as well as normal RAD51 recruitment at leptotene/zygotene stages (Suppl. Fig. 3c, d).

### Primary fibroblasts from BRC mutant mice display impaired IR-induced RAD51 recruitment

To compare the results observed in mESCs with somatic cells, we isolated embryonic fibroblasts (MEFs) of all genotypes and exposed them to IR to examine RAD51recruitment to DSBs. Unlike the observations made in mESCs, the homozygous and hemizygous mutant MEFs displayed significantly reduced number of RAD51 foci positive nuclei as well as reduced number of foci per nucleus, compared to control genotypes (Fig. 5a-c). To validate this result, we generated adult fibroblasts from ear punch and performed similar experiment. While the mutant adult fibroblasts showed an increase in RAD51 foci positive cells relative to the MEFs, these numbers remained significantly lower (Suppl. Fig. 4a, b).

**Fig. 5:**
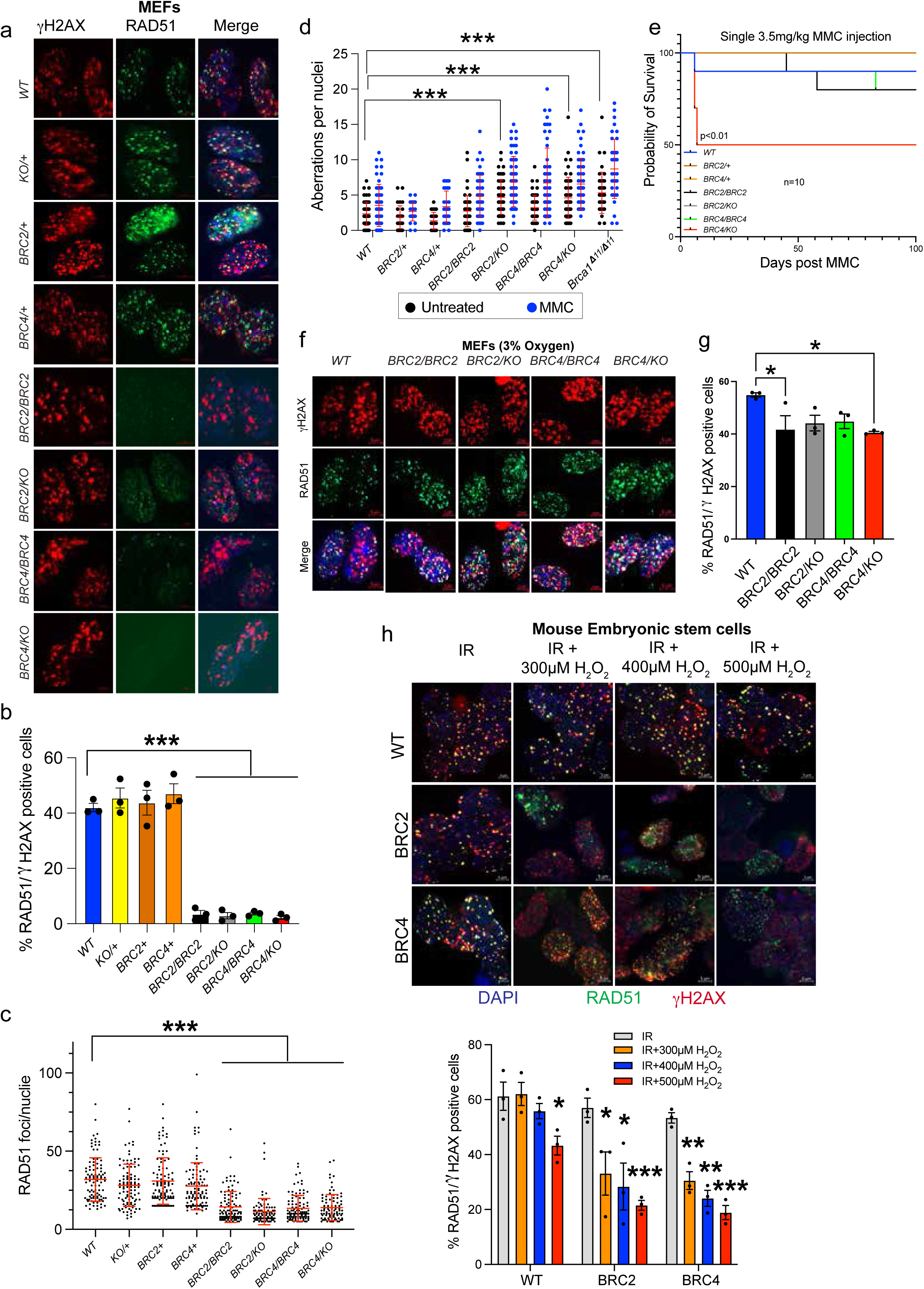
Primary fibroblasts from BRC mutant mice exhibit defect in RAD51 recruitment. a) Representative images depicting RAD51 foci at the DSBs generated by IR in MEFs isolated from mice of all genotypes. b) Quantification of percentage RAD51 positive nuclei observed in (a). Homozygous and hemizygous fibroblasts exhibited significantly lower number of RAD51 positive nuclei compared to WT (n=3, error bar-SEM, one way ANOVA, compared with WT). c) Quantification of RAD51 foci per nuclei in MEFs of all genotypes. Homozygous and hemizygous fibroblasts exhibited significantly lower number of RAD51 foci per nuclei (n=100, error bar-SD, one way ANOVA, compared with WT). d) Quantification of chromosomal aberrations per spread in MEFs of all genotypes in untreated and 100nM MMC treated conditions. *BRC2/KO* and *BRC4/KO* MEFs exhibited increased number of aberrations in untreated conditions as compared to WT (n=45, error bar-SD, one way ANOVA, compared with WT). MMC treatment exacerbates these aberrations in each genotype. *Brca1*^Δ11*/*Δ11^ MEFs are used as control for chromosomal aberrations. Representative images are shown in Suppl. fig. 4c. e) Mouse survival plot after single dose of MMC injections. *BRC4/KO* mice exhibited increased lethality and half of them dies within 3-weeks of injections (n=10, Log-rank Mantel–Cox test). f) Representative images depicting RAD51 foci at IR induced DSBs in MEFs of all genotypes cultured in low oxygen (3%). g) Quantification of percentage RAD51 positive nuclei observed in (f). Significantly increased percentages of RAD51 positive nuclei are visible in both homozygous and hemizygous fibroblasts compared to that in regular (21% oxygen) culture conditions (a, b) (n=3, error bar-SEM, one way ANOVA, compared with WT). h) Representative images depicting RAD51 foci at IR induced DSBs in mESCs under different concentrations of H_2_O_2_. i) Quantification of percentage RAD51 positive nuclei observed in (h). A dose dependent decline in percentages of RAD51 positive nuclei are visible in both BRC2 and BRC4 expressing mESCs (n=3, error bar-SEM, Students t-test, compared with IR). *p<0.05, p<0.01, ***p<0.001

A defect in RAD51 recruitment indicates a defect in DNA repair. To further assess this defect, we analyzed the MEFs for the presence of chromosomal aberrations with or without MMC treatment using *Brca1*Δ*11* (*Brca1*^Δ11*/*Δ11^) fibroblasts as control for genomic instability^26^. Interestingly, the hemizygous MEFs exhibited significantly higher chromosomal aberrations per nuclei compared to the WT MEFs in untreated condition (Fig. 5d, Suppl. Fig. 4c). However, all the mutant MEFs exhibited a significant increase in chromosomal aberrations after MMC treatment. The increased genomic instability in mutant MEFs further supports that BRCA2 containing BRC repeats 2 or 4 are not fully proficient in DNA repair.

Considering the defect in RAD51 recruitment in MEFs, we examined whether there was any difference in the protection of stalled RFs between mESCs and MEFs. We performed DNA fiber assay using *Brca1*^Δ11*/*Δ11^ fibroblasts, known to have a defect in RF protection^26^, as a control. Similar to the results observed in mESCs, mutant MEFs did not exhibit defect in stalled RF protection (Suppl. Fig. 4d).

Next, we assessed the sensitivity of the whole animal to DNA damage by injecting them with MMC. Studies show that mice with impaired HR die within three weeks of MMC injection^27^. Surprisingly, we observed that the MMC injection was well tolerated by mice of all genotypes except *BRC4/KO.* While half of *BRC4/KO* mice died within 3weeks of injection, the remaining survived until the end of the study (Fig. 5e). Such MMC sensitivity observed in *BRC4/KO* mice can be attributed to their lower body weight at the time of treatment (Fig. 4b).

The *in vivo* results suggest that BRCA2 with single BRC repeat is not fully functional contrary to results observed in mESCs expressing BRCA2 with BRC2 or BRC4 (Fig. 2). Especially the mutant mESCs were fully functional in RAD51 recruitment (Fig. 2b, c) as opposed to the mutant MEFs. We hypothesized that the differential culture conditions used to maintain mESCs and MEFs maybe responsible for the observed phenotype. mESCs are cultured in media consisting of β-mercaptoethanol (antioxidant), which provides a minimal oxidative stress^28^. However, fibroblasts are maintained in ambient oxygen levels (∼21%), without any antioxidant in the media, which is far greater than the *in vivo* physoxic environment (3-7.4% pO2)^29,30^. This difference in culture conditions may induce high oxidative stress in fibroblasts. To test this hypothesis, we cultured the MEFs under low oxygen (3%) conditions and examined IR-induced RAD51 recruitment. We observed a remarkable recovery in RAD51 recruitment in the mutant MEFs (Fig. 5f, g). In a complementary approach, we subjected the mESCs to oxidative stress by exposing them to varying concentrations of H_2_O_2_ along with IR. We observed a significant dose-dependent reduction in RAD51 recruitment in the mESCs expressing BRCA2 with single BRC repeats (BRC2/BRC4) (Fig. 5h, i). These results suggest that BRCA2 with a single BRC repeat is insufficient to cope with an additional oxidative stress, revealing the need to have multiple repeats.

### Residual HR has a dominant role relative to replication fork stability for mouse survival

Our previous study has shown that *Brca2^L2431P^* knock-in mice, expressing HR-deficient *Brca2* variant, do not survive in an *Mlh1*-null background^31^. *Mlh1*-null mice are defective in DNA mismatch repair as well as DNA2-mediated stalled RF protection^31^. We decided to examine the survival of our *BRC2* and *BRC4* mutant mice under this genetic stress by crossing them with *Mlh1* mutant mice (*Mlh1^-/+^*). Surprisingly, we observed survival of mice homozygous for *BRC2* and *BRC4* on *Mlh1^-/-^*background (*Brca2^BRC2/BRC2^;Mlh1^-/-^* and *Brca2^BRC4/BRC4^;Mlh1^-/-^*, referred as double mutant) in expected Mendelian ratios (Table 2). Additionally, primary fibroblasts from the ear punches were generated and examined for IR-induced RAD51 recruitment. Consistent with our previous results (Suppl. Fig. 4a, b), double mutant fibroblasts displayed reduced percentage of RAD51 foci positive nuclei compared to the controls (Fig. 6a, b). We also examined stalled RF protection in these fibroblasts and the double mutant fibroblasts displayed RF degradation after HU treatment as observed in *Mlh1^-/-^*fibroblasts (Fig. 6c). These findings confirm that residual HR is sufficient to support mouse viability even under defect in DNA mismatch repair and protection of stalled RFs.

**Fig. 6:**
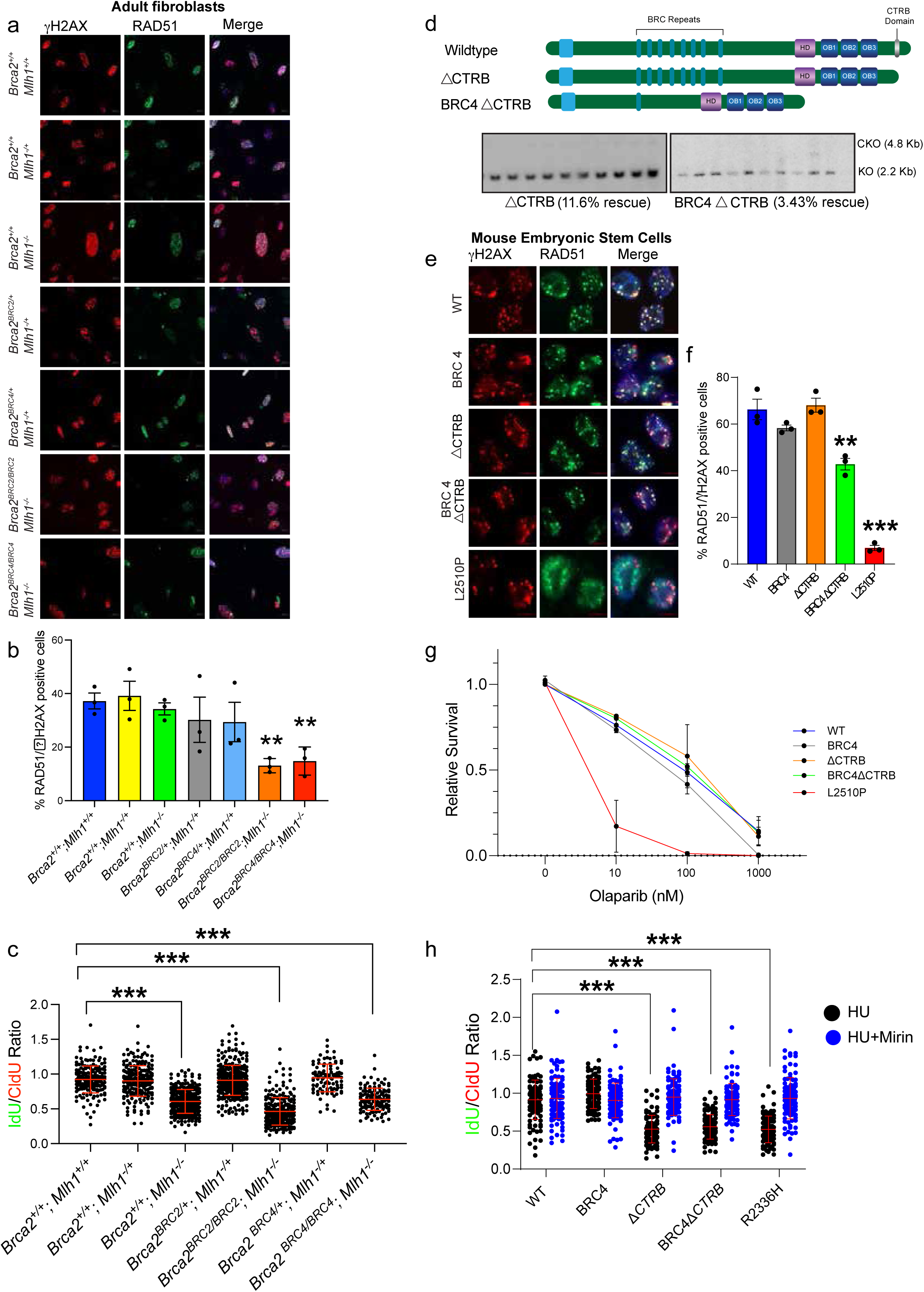
Replication fork defects do not hamper mouse and ES cell survival even with minimal HR. a) Representative images revealing RAD51 foci at the IR induced DSBs in adult fibroblasts isolated from mice of all genotypes in *Mlh1*-null background. b) Quantification of percentage RAD51 positive nuclei observed in (a). BRC2 and BRC4 homozygous fibroblasts (in *Mlh1*-null background) exhibited significantly lower number of RAD51 positive nuclei compared to WT (n=3, error bar-SEM, one way ANOVA, compared with WT). c) Replication fork stability measured by DNA fiber assay on fibroblasts of all genotypes in *Mlh1*-null background (explained in the schematic, Replication Fork is protected if the ratio of IdU/CldU tracks is close to 1, and it is degraded if the ratio is significantly reduced). *Mlh1*-null fibroblasts along with BRC2 and BRC4 homozygous fibroblasts (in *Mlh1*-null background) exhibited significant fork degradation as compared to WT (n>100, error bar-SD, Students t-test). d) Representative Southern blot images showing rescue percentage of ΔCTRB and BRC4ΔCTRB expressing mESCs. e) Representative images depicting RAD51 foci at DSBs generated by IR in ΔCTRB and BRC4ΔCTRB expressing mESCs. f) Quantification of percentage RAD51 positive nuclei observed in (e). BRC4ΔCTRB expressing mESCs exhibited significantly lower number of RAD51 positive nuclei compared to WT (n=3, error bar-SEM, one way ANOVA, compared with WT). g) Colony forming assay to determine olaparib sensitivity of mESCs expressing ΔCTRB and BRC4ΔCTRB. Only the known hypomorphic BRCA2 variant L2510P mESCs exhibited significant sensitivity towards olaparib (n=3, error bar-SEM, Students t-test, two tailed). h) Replication fork stability measured by DNA fiber assay on ΔCTRB and BRC4ΔCTRB expressing mESCs (in presence of HU and HU+Mirin). ΔCTRB and BRC4ΔCTRB expressing mESCs exhibited significant fork degradation as compared to WT which was rescued in presence of MRE11 inhibitor Mirin (fork protection when IdU/CldU ≥1, n>100, error bar-SD, Students t-test). **p<0.01, ***p<0.001

**Table 2.** Mating of *BRC2* and *BRC4* mutant mice with *Mlh1* mutant mice.

| *Brca2<sup>BRC2/+</sup>;Mlh1<sup>-/+</sup>* X<br>*Brca2<sup>BRC2/+</sup>;Mlh1<sup>-/+</sup>* | Observed | Expected | *Brca2<sup>BRC4/+</sup>;Mlh1<sup>-/+</sup>* X<br>*Brca2<sup>BRC4/+</sup>;Mlh1<sup>-/+</sup>* | Observed | Expected |
| --- | --- | --- | --- | --- | --- |
| *Brca2<sup>+/+</sup>;Mlh1<sup>+/+</sup>* | 10 | 13 | *Brca2<sup>+/+</sup>;Mlh1<sup>+/+</sup>* | 14 | 8.8125 |
| *Brca2<sup>BRC2/+</sup>;Mlh1<sup>+/+</sup>* | 44 | 26 | *Brca2<sup>BRC4/+</sup>;Mlh1<sup>+/+</sup>* | 26 | 17.625 |
| *Brca2<sup>BRC2/+</sup>;Mlh1<sup>-/+</sup>* | 40 | 52 | *Brca2<sup>BRC4/+</sup>;Mlh1<sup>-/+</sup>* | 30 | 35.25 |
| *Brca2<sup>BRC2/BRC2</sup>;Mlh1<sup>+/+</sup>* | 19 | 13 | *Brca2<sup>BRC4/BRC4</sup>;Mlh1<sup>+/+</sup>* | 16 | 8.8125 |
| *Brca2<sup>BRC2/BRC2</sup>;Mlh1<sup>-/+</sup>* | 26 | 26 | *Brca2<sup>BRC4/BRC4</sup>;Mlh1<sup>-/+</sup>* | 14 | 17.625 |
| *Brca2<sup>BRC2/BRC2</sup>;Mlh1<sup>-/-</sup>* | 12 | 13 | *Brca2<sup>BRC4/BRC4</sup>;Mlh1<sup>-/-</sup>* | 8 | 8.8125 |
| *Brca2<sup>+/+</sup>;Mlh1<sup>-/+</sup>* | 29 | 26 | *Brca2<sup>+/+</sup>;Mlh1<sup>-/+</sup>* | 14 | 17.625 |
| *Brca2<sup>+/+</sup>;Mlh1<sup>-/-</sup>* | 9 | 13 | *Brca2<sup>+/+</sup>;Mlh1<sup>-/-</sup>* | 8 | 8.8125 |
| *Brca2<sup>BRC2/+</sup>;Mlh1<sup>-/-</sup>* | 19 | 26 | *Brca2<sup>BRC4/+</sup>;Mlh1<sup>-/-</sup>* | 11 | 17.625 |
| $\chi^2$ value | P=0.004 | | $\chi^2$ value | P=0.022 | |

The critical function of BRCA2 in protecting MRE11-mediated stalled RF degradation is by its CTRB (residues 3260-3314) domain encoded by exon27^21^. Moreover, a study reported CTRB is required for the RAD51 nucleation at the DSBs^32^. To ascertain which of the two RAD51 binding motifs, the BRC repeats or the CTRB, is critical we deleted the CTRB domain in a BAC containing full length BRCA2 (ΔCTRB) and BRCA2 with BRC4 (BRC4ΔCTRB). Expressing these BACs in our mESCs revealed higher rescue rate (11.5%) in ΔCTRB as compared to that in BRC4ΔCTRB (3.43%) (Fig. 6d). Interestingly, the BRC4ΔCTRB rescue rate was comparable to the BRCA2 with BRC4 alone (Fig. 2a). Furthermore, mESCs expressing ΔCTRB exhibited comparable ability of RAD51 recruitment with WT and BRC4 cells. However, a slight reduction in RAD51 recruitment was observed in cells expressing BRC4ΔCTRB (Fig. 6e, f). Next, we treated these cells with olaparib and found no difference in the sensitivity of mESCs expressing ΔCTRB or BRC4ΔCTRB compared to WT BRCA2 (Fig. 6g). Furthermore, examining the stalled RF protection ability resulted in consistent fork degradation in cells expressing either ΔCTRB or BRC4ΔCTRB, which was suppressed in the presence of mirin (MRE11 inhibitor) (Fig. 6h). These findings unequivocally demonstrate the division of functions among the two RAD51 binding domains of BRCA2. Furthermore, the failure to obtain viable mESCs or mice upon deletion of all BRC repeats demonstrates that BRC repeats are critical for survival in contrast to the CTRB domain.

## Discussion

BRCA2 functions as a tumor suppressor because of its role in error free DNA repair by HR^1^. It recruits RAD51 to the DSBs, via the eight BRC repeats in the middle of the protein and the CTRB near the C-terminal end^33^. Most species have multiple BRC repeats, including mice and humans (eight BRC repeats), and several previous studies have revealed functional redundancy in them^5,16,33^. Yet, the functional significance of retaining multiple repeats remains largely unknown. Our study addresses this long-standing question by examining, for the first time *in vivo*, the requirement for multiple BRC repeats within BRCA2. By using humanized mESCs, expressing *BRCA2*, we observed the BRC repeats 5-8 were dispensable for specific BRCA2 functions. We identified the most crucial BRC repeat for BRCA2 function and confirmed that BRCA2 harboring a single BRC repeat (BRC2 or BRC4) is fully functional in the mESCs, albeit with reduced HR efficiency.

We failed to obtain viable mice lacking all the BRC repeats in hemizygous or homozygous states confirming our *in vitro* results. However, knock-in mouse models with BRCA2 harboring a single BRC repeat (BRC2 or BRC4) in homozygous as well as hemizygous state are viable and do not show any overt phenotype. This suggests that approximately one third of BRCA2 is dispensable for survival. Additionally, BRC2 mutant mice performed better across several developmental parameters compared to BRC4 mutants, suggesting superior *in vivo* performance of BRC2 relative to BRC4. We observed reduced HR efficiency in mESCs expressing BRCA2 with BRC2 or BRC4, but this did not hamper the BRC mutant mice survival in *Mlh1*-null background, defective in DNA mismatch repair and replication fork protection.

We were puzzled by the severe defect in RAD51 recruitment and increased chromosomal aberrations in the mutant MEFs. This was in sharp contrast to the efficient RAD51 recruitment observed in mESCs and spermatocytes (during meiosis). We hypothesized that the inconsistencies observed in the RAD51 recruitment abilities between fibroblasts and mESCs can be attributed to the different culture conditions of these cell lines. Regular culture conditions expose MEFs to ambient oxygen (21%), which is much higher than the 3-7.4% pO_2_ in tissues inducing oxidative stress^29,30^. However, mESCs are cultured in the presence of a feeder layer (along with β-mercaptoethanol in the media) that provides a highly supportive microenvironment ensuring pluripotency in them under minimal oxidative stress^28^. Remarkably, the mutant MEFs exhibited a significant increase in RAD51 recruitment when cultured in low oxygen (3%) conditions. Likewise, we observed a decline in RAD51 recruitment ability of mESCs expressing BRCA2 with BRC2 or BRC4 in presence of H_2_O_2_. These results along with the reduced binding of RAD51 with single BRC repeat containing BRCA2 (Fig. 2e,f), suggest that the oxidative stress might further destabilize this interaction and the presence of multiple BRC repeats provide robustness. Similar phenomena can also explain the reduction in mutant HSPC colonies and their hypersensitivity to olaparib, yet the mice were not sensitive to proliferative stress induced by 5FU.

In conclusion, our functional studies demonstrate that BRC repeats are essential for the survival of cells and mice. Despite the presence of eight BRC repeats in human and mouse BRCA2, a single BRC repeat is sufficient for RAD51 recruitment to DSBs even in the absence of the CTRB domain. We have shown that 1093 amino acids (out of 3329) of mouse BRCA2, containing 7 BRC repeats, are dispensable for its critical function in HR and RF protection. We conclude that the residual HR present in the mutant (homozygous and hemizygous) mice is sufficient for their physiological growth and development but inefficient when the mutant cells are subjected to oxidative stress. We predict that inducing oxidative stress may resensitize certain HR-proficient cancers, with large in-frame deletion in exon11 retaining few BRC repeats, that are resistant to olaparib^15,34^.

## Methods

### Generation of mESCs harboring BRCA2 with different deleted BRC repeats

Different BRC repeat deletions in *BRCA2* were generated using *BRCA2* cloned in a Bacterial artificial chromosome (BAC). We used the oligonucleotide-based ’hit and fix’ BAC recombineering based approach as described earlier^35^. Correctly targeted bacterial clones were confirmed by PCR and sequencing. We electroporated 25μg BAC DNA into 10^7^ actively dividing Pl2F7 mESCs and selected the colonies with geneticin (180μg/ml, 10131-035, Gibco). Geneticin resistant colonies were picked on a 96-well plate, and the total RNA was isolated. Expression of the integrated BAC was confirmed by one step reverse transcriptase PCR [G597, ABM) primer sequence in Extended Table 1]. The RT-PCR positive colonies were expanded and subjected to Cre-HAT selection as described earlier^17^. Colonies were picked in 96-well plate and genomic DNA was isolated. Surviving colonies were identified using Southern blot analysis on these colonies as explained earlier^17^. The percentage of rescued colonies (showing only the KO band on the southern blot, Suppl. Fig. 1B) was multiplied by the proportion of colonies acquired on a HAT plate as opposed to that on a no-HAT plate to get the rescue percentage.

### Assay to test sensitivity for different DNA damaging drugs

Actively dividing mESCs were plated in 96-well plates (10,000 cells/well). Required concentrations of DNA damaging drugs (cisplatin, MMC, olaparib and IR) were added to each well and incubated at 37°C. After 72hrs the plates were washed with PBS and 100μl of XTT solution (J61726, Sigma) solution (1mg/ml in phenol red free DMEM, 21041025, Gibco) was added. After incubation at 37°C for one hour the colorimetric readings were taken and the graph was plotted on GraphPad Prism.

### IR-induced RAD51 foci formation

Cells were seeded (10^5^ cells /well) on poly-d-Lysine coverslips (GG-12, Neuvitro) and irradiated 6Gy for mESCs and 10Gy for fibroblasts). After recovery (6hrs for mESCs and 3hrs for fibroblasts) the cells were treated with a hypotonic solution (85.5 mM NaCl, 5 mM MgCl2, pH 7) for 10mins and then fixed (4% paraformaldehyde and 0.03% SDS in PBS) for 10mins at room temperature. Cells were incubated overnight at 4°C with primary antibodies [γH2AX (1:500, JBW301, Millipore), RAD51 (1:250, PC130, Millipore)] diluted in antibody dilution buffer (1% BSA, 0.3% TritonX100, 5% goat serum in PBS). Next morning, the cells were washed 3X with PBST (PBS containing 0.2% Triton X-100) and incubated with secondary antibodies [Alexa-fluor anti-mouse 594 (1:1000, A11005, Invitrogen) and anti-rabbit 488 (1:500, A11034, Invitrogen)] diluted in PBS at 37°C for one hour. The cells were washed 3X with PBST and stained for one minute with DAPI (1:50,000, 11190301, Sigma). Clean, labeled slides were used to mount the coverslips using anti-fade mount (P36930, Invitrogen). The slides are examined using Zeiss AXIO imager M2 (63X).

### RAD51 foci formation in mESCs under oxidative stress

mESCs expressing different BRCA2 constructs were cultures in M15 media. 105 cells were plated on coverslips. Next day the media was changed to varying doses of H2O2 dissolved in non β-mercaptoethanol containing DMEM media. After 3 hrs, these were given 6GY of IR and the media was changed to fresh H2O2 containing DMEM media. After 6 hrs, cells were stained for RAD51 and γ-H2AX as described above.

### HR assay

We used mESCs (F7A10) with 29 bp deleted blasticidin resistance gene^19^. 2×10^6^ actively dividing F7A10 cells (harboring WT BRCA2 or with different BRC repeats) were plated. After 24 h cells were nucleofected using nucleofector kit (VPH-1001, Lonza) with 2 µg of gRNA-Cas9 expressing plasmid px330 (Addgene plasmid #158973), and 1 µg of linearized donor blasticidin sequence (supplemented with the 29bp, PAM mutated) cloned in Topo-TA vector (Invitrogen 450641) as per manufacturer’s guidelines. Cells were resuspended in 5 ml M15 media (Knockout DMEM, 15% FBS, 1X β-mercaptoethanol) and plated on 60 mm culture dish with blasticidin-resistant feeders. 48hrs post plating the cells were treated with blasticidin (15 µg/ml in M15 media, 11139-03 Gibco) for five days. After the treatment, cells were incubated in M15 media until the colonies started to appear. Assessment of HR was performed by staining the plates with methylene blue (0.05% in 70% ethanol) and colonies were counted.

### Amylose Pull-downs

Transient transfection of 1µg of each construct (phCMV1 mammalian expression vector including a 2XMBP fusion to BRCA2 containing the BRC repeat(s) indicated) into actively dividing 5×10^5^ 293T cells was done in 6-well plates using TurboFect (R0531, ThermoFisher Scientific). After 36hrs the cells were harvested in 500µL of the lysis buffer [50mM HEPES (pH7.5), 250mM NaCl, 1% Igepal CA-630, 1mM MgCl2, 1mM DTT, 250 Units/mL Benzonase (EMD Millipore), and 1X EDTA-free protease inhibitor cocktail (C762Q72, Roche)]. Total cellular lysate aliquots were taken before batch binding for protein expression analysis. Cell lysates were batch bound to 20µL of amylose resin for 2hrs to capture the 2XMBP tagged BRCA2 proteins, washed 3X in wash buffer [50mM HEPES (pH7.5), 250mM NaCl, 0.5mM EDTA, and 1mM DTT]. Proteins were then eluted in the same buffer containing 10mM maltose and 10% glycerol. Samples were run on 4-15% gradient SDS-PAGE TGX stain-free gels (456-8086, Bio-Rad). The 2XMBP-BRCA2 proteins (amylose pull-downs) were visualized by Stain-Free imaging on a ChemiDoc MP imaging system (Bio-Rad). Total cell lysate gels were also visualized by Stain-Free imaging to ensure equal loading. BRCA2 and RAD51 proteins were detected by transferring gels to PVDF (IPVH00010, Millipore) membranes overnight at 4°C, blocking for 30 minutes with 5% milk in 1XTBS-T [50 mM Tris (pH7.5), 150 mM NaCl, 0.05% Tween20], incubating the membranes overnight with either anti-MBP (1:5000, E8032L, NEB) or anti-RAD51 antibody (1:1000, PC130 Millipore) in 1XTBS-T. Membranes were washed 3 times with TBS-T and then incubated with secondary mouse and rabbit antibodies (HRP-conjugated, sc-516102 and sc-2004, respectively, Santa Cruz Biotechnology). The Western blots were visualized using Clarity Western ECL substrate (170-5061, Bio-Rad) for five minutes and visualized on a ChemiDocMP system. Band densitometry was performed using ImageLab (Bio-Rad).

### DNA fiber assay

In a 6-well plate, 5 × 10^5^ cells were plated and treated with thymidine analogues 8μg/ml CldU (8µg/ml) and IdU (90μg/ml) for 30 minutes each, followed by a 4-hour treatment with 4 mM hydroxyurea (HU). After treatments, the cells were trypsinized and resuspended in PBS. Cell suspension and lysis buffer were added to the slide to accomplish cell lysis. The slides were tilted to allow the fibers to spread and air dry after incubation for roughly ten minutes. The fibers were fixed overnight using a methanol:acetic acid (3:1) mixture, then rehydrated using PBS and denatured for one hour in 2.5M HCl. After washing with PBS, the slides are blocked for 40 minutes using 5% BSA. The primary mouse anti-BrdU antibody (1:500, 347580, Becton Dickinson) and the rat anti-BrdU antibody (1:500, ab6326, Abcam) were incubated on the fibers for two hours after blocking. After rinsing with PBST, the slides were treated for one hour at room temperature with secondary anti-mouse AlexaFluor488 (1:500, A21202, Invitrogen) and anti-rat AlexaFluor594 (1:500, A11007, Invitrogen). The slides were washed three times with PBST and were mounted (P36930, Invitrogen) and examined using Zeiss AXIO imager M2 (63X). Green and red fiber lengths (at least 100 fibers per sample) were measured using Fiji software, and their ratios were calculated and shown^36^.

### Saturation Genome Editing in BRC repeats

We used an mESC line expressing a single copy of human *BRCA2* (*Brca2^−/−^; Tg[BRCA2]*) [Clone: F7/F7]^24^. Two sgRNAs targeting regions upstream of the BRC1 repeat and downstream of the BRC8 repeat were cloned into the pX458-Cas9-GFP plasmid. Three million cells were nucleofected with these plasmids and a ssODN containing only the BRC2 repeat using the Lonza Nucleofector 2B (program A030). GFP-positive cells were sorted and plated, and 96 colonies screened to isolate a clonal line containing only the BRC2 repeat (*Brca2^−/−^; Tg[BRCA2^BRC2^]*). This line was used for subsequent saturation genome editing (SGE) experiments, where oligo donor pools containing either individual BRC repeats or NNN degenerate codons for all possible amino acid substitutions at defined residues were nucleofected into mESCs. Post-nucleofection, cells were pooled, treated with cisplatin, olaparib, or DMSO, and harvested for DNA extraction and deep sequencing, with data analyzed as previously described^23^. BRC repeats counts and frequencies were extracted from FASTQ files using a custom Python script (v 3.10.12). Heatmaps were generated in R Studio (v4.4.0) using the ggplot2 (v 4.0.1) and ComplexHeatmap packages (2.20.0)^37,38^.

### Generation of *Brca2^BRC2/+^* and *Brca2^BRC4/+^* Knock-in mice

Fertilized zygotes (0.5dpc) were isolated from C57Bl/6 female mouse. These zygotes were microinjected with ribonucleoprotein complex (RNP complex) containing pure Cas9 protein, upstream and downstream gRNAs for exon11, and donor DNA (containing either for BRC2 or BRC4) (for sequences see supporting document). As explained in Suppl. Fig. 2A, both the gRNAs help in excising out all the eight BRC repeats from exon11 from one of the *Brca2* alleles and are repaired by homologous recombination using the flanking sequences of the donor DNA provided. The injected zygotes were implanted in pseudo-pregnant females for embryo development. Live pups obtained from these females are weaned at 3 weeks of age and tail clips are obtained for genotyping and sequencing (Suppl. Fig. 2b,c, Suppl. Table 1). Sequencing confirmed animals are back crossed to C57Bl/6NCR mice (for at least 10 generations) and maintained as separate mouse lines of BRC2 and BRC4.

### Ethics statement

The Guide for the Care and usage of Laboratory Animals (The National Academies Press; 8th edition) was followed in the housing, breeding, and study usage of all mice. The NCI-Frederick Animal Care and Usage Committee (ACUC) approved the study protocol (Animal Study# 24-471). The animals were kept in a 12-hour cycle of light and dark. Temperatures between 20°C-27°C and relative humidity levels between 30 and 70% were maintained in the rooms. All animal studies were performed in compliance with ARRIVE (Animal Research: Reporting In Vivo Experiments) guidelines (https://arriveguidelines.org/arrive-guidelines).

### Mammary gland isolation and Carmine Alum staining

Five-week-old female mice were used to harvest their mammary glands, which were then preserved in Carnoy’s solution (6:3:1 of ethanol, chloroform, and glacial acetic acid). Carmine Alum (C1022, Sigma) was used to stain the glands overnight after they had been rehydrated using progressively lower grades of ethanol wash. After being dried in progressively higher grades of ethanol, the glands were incubated for 2-3 days in xylene replacement to clean the tissue. The whole-mount glands were evaluated after they were photographed with a Zeiss Axiocam brightfield microscope.

### Colony forming assay from fetal liver cells

Embryos were collected at E16.5 from pregnant females and euthanized. These embryos’ livers were meticulously removed and tweezed in IMDM. A 40µm cell strainer was used to filter the cell suspension. MethoCult (M3231, Stem Cell Technologies) media with growth factors (10% FBS, 100ng/ml muSCF, 100ng/ml huTpo, 100ng/ml huFlt3L, 50ng/ml muIL6, 30ng/ml muIL3) were used to plate 25,000 live cells. These were incubated at 37°C for 7-10 days. Iodonitrotetrazolium chloride (1mg/ml, I10406, Sigma) was used to stain the colonies.

### Exhaustion of hematopoietic cells by multiple 5-Flurouracil injection

Six to eight weeks old mice from each genotype were intraperitoneally injected (weekly) with 5-Flurouracil (5FU) (135mg/kg, NDC68001-525-27, USP grade, BluePoint Laboratories) for 3 weeks. After 50 days from first injection, the animals were again injected with 5FU continuously every week until they started to show mortality. In accordance with ACUC procedures, mice exhibiting symptoms of distress, such as weight loss, were euthanized.

### Generation of embryonic mouse fibroblasts

E13.5 embryos were obtained from timed mating of different genotypes. After genotyping the embryos, they were minced in 0.5% trypsin (15400-054, Gibco) and incubated at 37°C for 30 minutes. After incubation the slurry was plated on 100mm cell-culture plates in DMEM+10% FBS in 37°C, 5% CO_2_ incubator. After they became confluent, small aliquots of mouse embryonic fibroblasts (MEFs) were frozen in liquid nitrogen as P0 (passage zero).

### Generation of adult fibroblasts

Ear punch from the animals of required genotype were obtained. After a brief wash in 70% ethanol, they were rinsed in sterile Hank’s balanced salt solution (HBSS). The ear punches were finely minced in collagenase solution (2000U/ml in HBSS, C7657, Sigma) and incubated at room temperature (on a roller mixer) for 3hr. Digested ear punches were centrifuged, and supernatant was removed. Rest of the procedure is same as discussed above for embryonic fibroblasts.

### Analyses of chromosomal aberrations

MEFs of desired genotypes were incubated with MMC (100nM, 11435, Cayman Chemicals) for 12hr and released in normal media for 12hr. After release, the cells were arrested in metaphase using colcemid (10ug/ml, 15210-016, KaryoMAX) dissolved in DMEM and incubation for 12hr. The cells were fixed in methanol: acetic acid (3:1) and metaphase spreads were made on a clean labeled slide. Slides were stained with Giemsa, and the chromosomal aberrations were quantified.

### Animal survival study post MMC injection

Six to eight weeks old animals from each genotype were intraperitoneally injected with MMC (3.5mg/kg, single dose, NDC 55150-451-01, USP grade, Eugia). Animals were observed every day till they started to show mortality. In accordance with ACUC procedures, mice exhibiting symptoms of distress, such as weight loss, were euthanized.

### Histology

Tissues (ovaries and testes) were overnight fixed in 10% formalin, paraffin-embedded, sectioned (5µm) and stained for H&E (hematoxylin and eosin). The stained slides were visualized under bright field microscope.

### Meiotic chromosomes spread and analysis

Male mice aged 4-6 weeks had their testes used to prepare meiotic spreads. The testes were placed in hypo-extraction buffer PBS (30 mM Tris pH8.2, 50 mM sucrose, 17 mM citric acid, 5 mM EDTA, 0.5 mM DTT, and 0.1 mM PMSF) after being quickly rinsed in. After carefully removing the tunica from the testes and removing the tubules with fine forceps, the mixture was incubated at room temperature for half an hour. 25µl of 0.1M sucrose solution and a tiny piece of the digested tissue segment from the previous stage were put on a clean labeled slide that had been prerinsed with PFA (4%, pH 9.2 adjusted with 50 mM boric acid). This tissue was shredded and spread out on the slide using a fine needle. For slow drying the slides were kept in a humid chamber overnight. The immunofluorescence staining was performed using primary antibodies: mouse anti-SYCP3 (1:500, sc-74568, Santa Cruz); rabbit anti-RAD51 (1:250, PC130, Millipore). Secondary antibody staining and further processes were performed as described in the RAD51 foci formation assay section.

### Generation of *Brca2^BRC2^* and *Brca2^BRC4^* animals in *Mlh1^-/-^* background

We mated the *Brca2^BRC2/BRC2^* and *Brca2^BRC4/BRC4^*mice with *Mlh1^-/+^* mice (*Mlh1^-/-^* mice are infertile)^39^ to obtain heterozygous mice for both the genes (*Brca2^BRC2/+^;Mlh1^-/+^* and *Brca2^BRC4/+^;Mlh1^-/+^*). We intercrossed these double heterozygous animals to obtain double mutants (*Brca2^BRC2/BRC2^;Mlh1^-/-^*and *Brca2^BRC4/BRC4^;Mlh1^-/-^*).

### Statistical analysis

Microsoft Excel and GraphPad Prism version 6.0 were used for all statistical tests. Figure legends for all experiments explain the exact statistical test and p-values along with the error bars.

### Biological material availability

All unique materials used in the study are readily available from the authors upon request as indicated in the Methods section.

## Data availability

All relevant data are provided as supplementary information. Any additional information will be provided upon request.

## Code availability

https://github.com/MelissaGall/BRC_saturation_heatmap

## Acknowledgement

We thank Drs. Ira Daar and Jonathan Keller for helpful discussions and critical review of the manuscript. This research was supported in part by the Intramural Research Program of the National Institutes of Health (NIH) (L.T. and S.K. Sharan). The contributions of the NIH author(s) were made as part of their official duties as NIH federal employees, are in compliance with agency policy requirements, and are considered Works of the United States Government. However, the findings and conclusions presented in this paper are those of the author(s) and do not necessarily reflect the views of the NIH or the U.S. Department of Health and Human Services. R.B.J. was supported by an NIH grant (R01 CA270788).

## Author Contribution

APM conceived, designed and performed most of the experiments and analyzed the results; SS, ES, TS, DC performed CRISPR/cas9-based SGE; GM, JRJ, RBJ performed in vitro RAD51 binding assay; SKSengodan performed DNA fiber assays, NO generated BRC mutant BAC constructs, SP performed mammary gland studies, MG performed the bioinformatics analysis; M.A. helped with mice studies. PPA generated the knock-in mice, FTA and LT helped with gRNAs and donor DNA design for knock-in mice. SB performed cytogenetic analysis, APM and SS prepared the figures; SKSharan conceived and supervised the study. APM and SKSharan wrote the manuscript, and all authors reviewed and edited the manuscript.

