## Supplementary Figures 1-4 and Suppl Table 1 for "A single BRCA2 BRC repeat supports viability while multiple repeats ensure resilience under stress"

**Supplementary information**

**Supplementary Fig. 1** a) Graphical representation of number of BRC repeats present in BRCA2 homologues in different species in increasing order of complexity. Notably, different species in *Trypanosoma* genus have varied number of BRC repeats. b) Schematic of strategy to genotype HAT rescued mESCs after mutant *BRCA2* reconstitution in PL2F7 cells genomic DNA digestion using *EcoRV* and Southern blot analysis. c, d) Representative Southern blot images showing rescue percentage of mESCs expressing BRCA2 with different number of BRC repeats intact. mESCs with all the BRC repeats deleted (BRCΔ1-8) does not reveal any rescued clone showing both the CKO and KO bands in all the lanes. e) Replication fork stability measured by DNA fiber assay in mESCs expressing BRCA2 with different number of BRC repeats (explained in the schematic, replication fork is protected if the ratio of IdU/CldU tracks is close to 1, and it is degraded if the ratio is significantly reduced). Only the known BRCA2 hypomorph R2336H exhibited significant fork degradation as compared to WT (n>100, error bar-SD, Students t-test).

### Supplementary figure 1

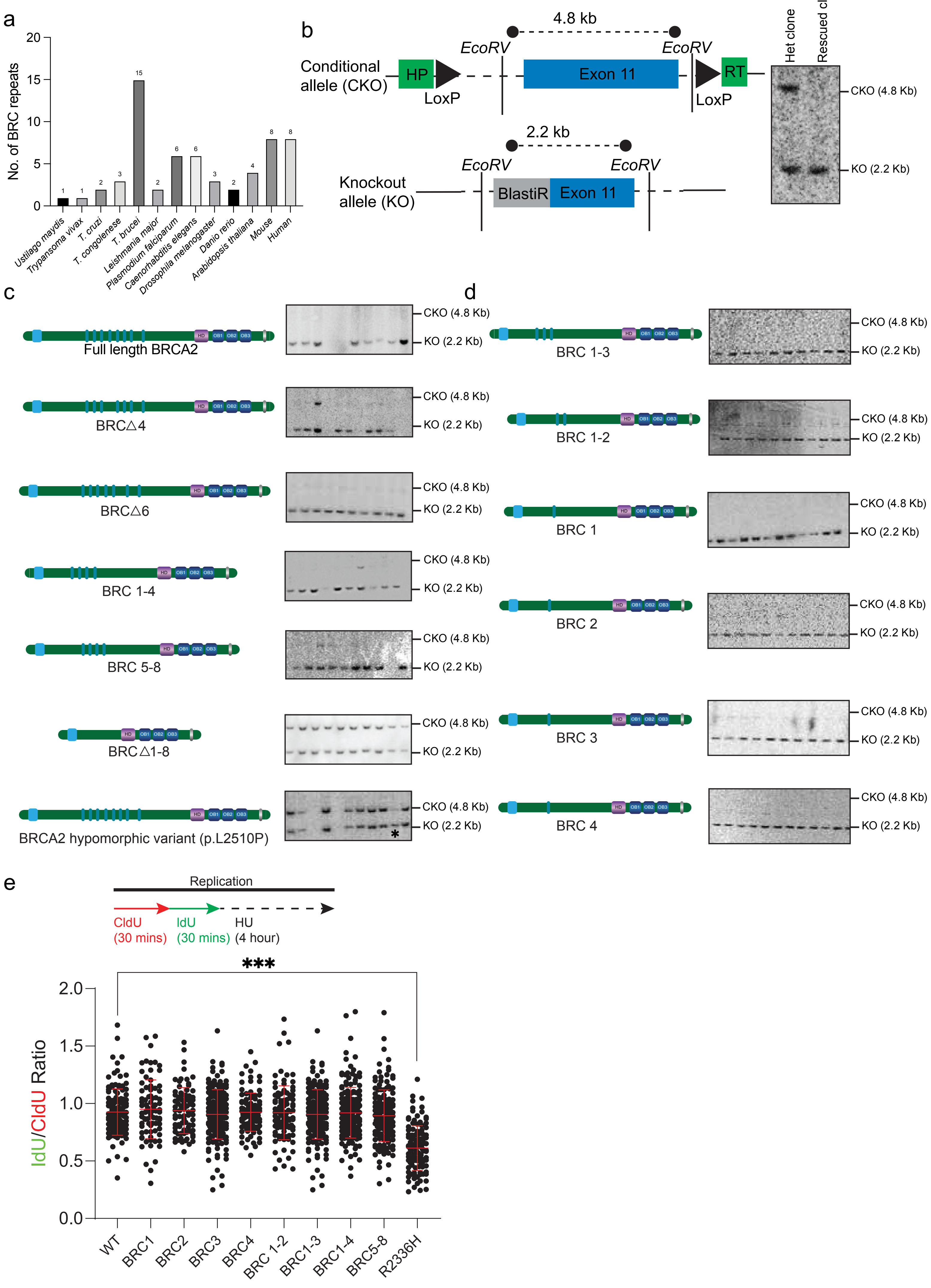

**Supplementary Fig. 2** a) Schematic of generation of knock-in mouse model for single BRC repeat of BRCA2. RNP complex, containing Cas9 protein, sgRNA1&2 and ssODN of BRC2 or 4, is microinjected in the 0.5dpc zygote and transplanted in pseudo-pregnant females for embryos to develop and deliver. The resultant size of BRCA2 is one-third smaller than the full length. Genomic loci drawn not to scale. b) Genotyping strategy of pups obtained from the above-mentioned technique. c) Confirmation of rightly targeted mice for BRC2 and BRC4 by sequencing. One animal with in-frame deletion of all the BRC repeats (BRCΔ1-8) was also obtained as confirmed by sequencing.

Supplementary figure 2

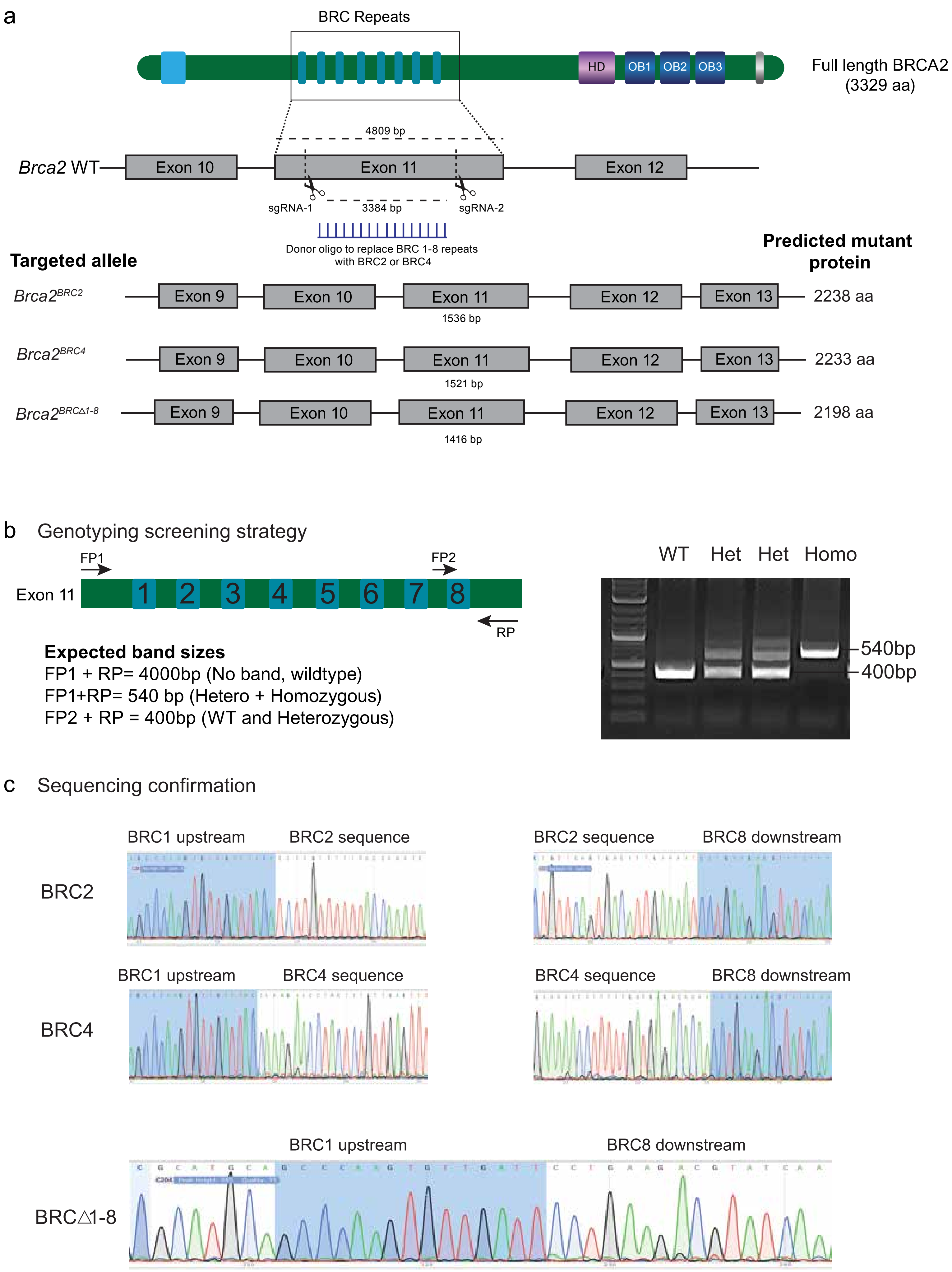

**Supplementary Fig. 3** a) Representative images showing CFUs from fetal liver cells from different genotypes with and without 100nM olaparib treatment. b) H&E analyses of testes and ovaries from WT and mutant animals (3-4 weeks old) show normal histology with sperm in seminiferous tubules and regular follicular development in ovaries. c) Spermatocyte spreads from tetes of all genotypes at Leptotene/Zygotene stages. Homologous chromosomes are labeled with SYCP1, and RAD51 foci formation is witnessed. d) Quantification of number of RAD51 foci present per spermatocyte spread for all genotypes (n=100). All genotypes exhibited comparable number of RAD51 foci per spermatocyte spread.

Supplementary figure 3

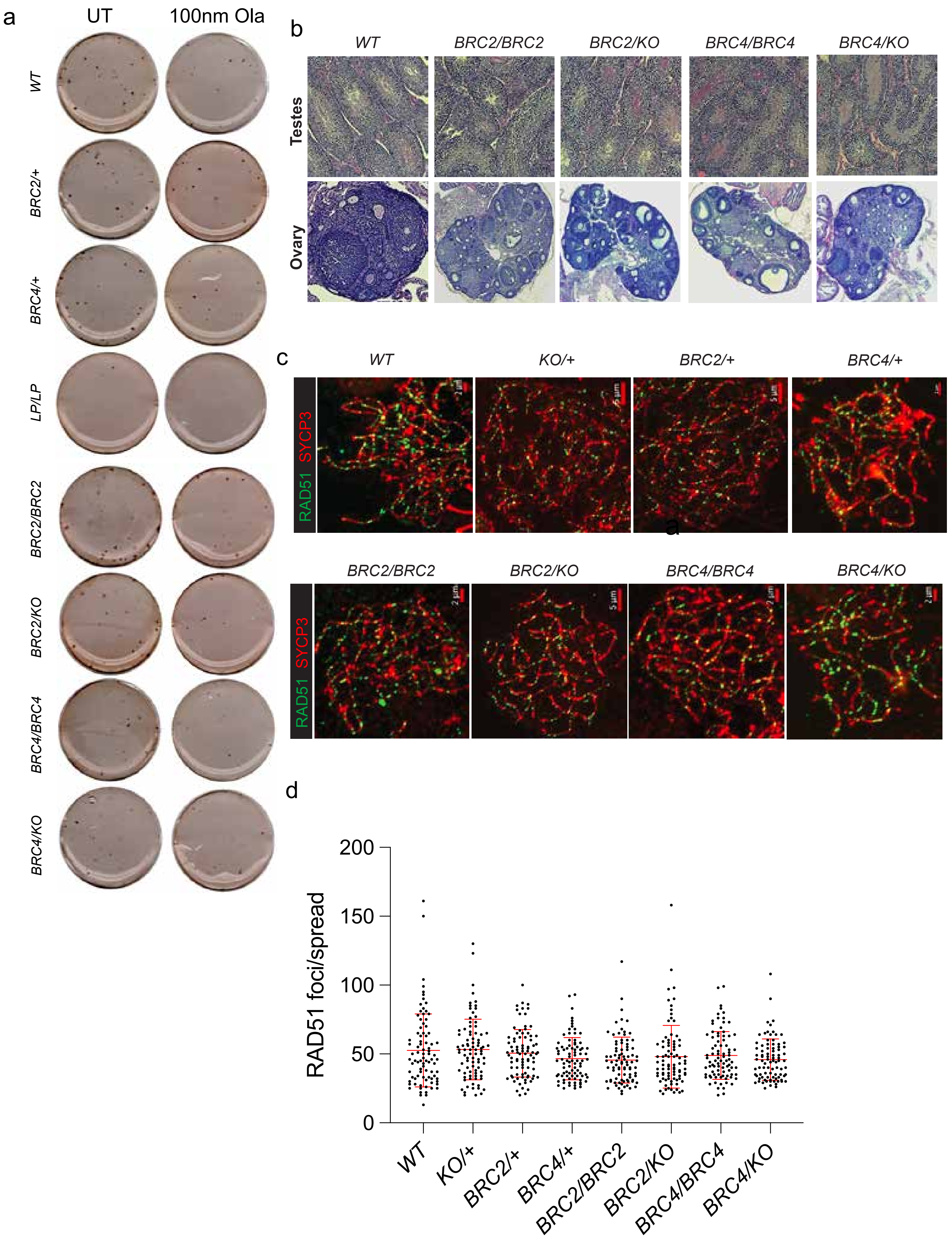

**Supplementary Fig. 4** a) Representative images depicting RAD51 recruitment at IR-induced DSBs in adult fibroblasts of all genotypes. b) Quantification of percentage RAD51 positive nuclei observed in (c). Homozygous and hemizygous fibroblasts exhibited significantly lower number of RAD51 foci per nuclei (one way ANOVA, compared with WT). c) Representative images of chromosomal aberrations in MEFs of all genotypes in untreated and 100nM MMC treated conditions. d) Replication fork stability measured by DNA fiber assay on MEFs of all genotypes (explained in the schematic). Only *Brca1Δ11* MEFs exhibited significant fork degradation as compared to WT (n>100, error bar-SD, Students t-test).

### Supplementary figure 4

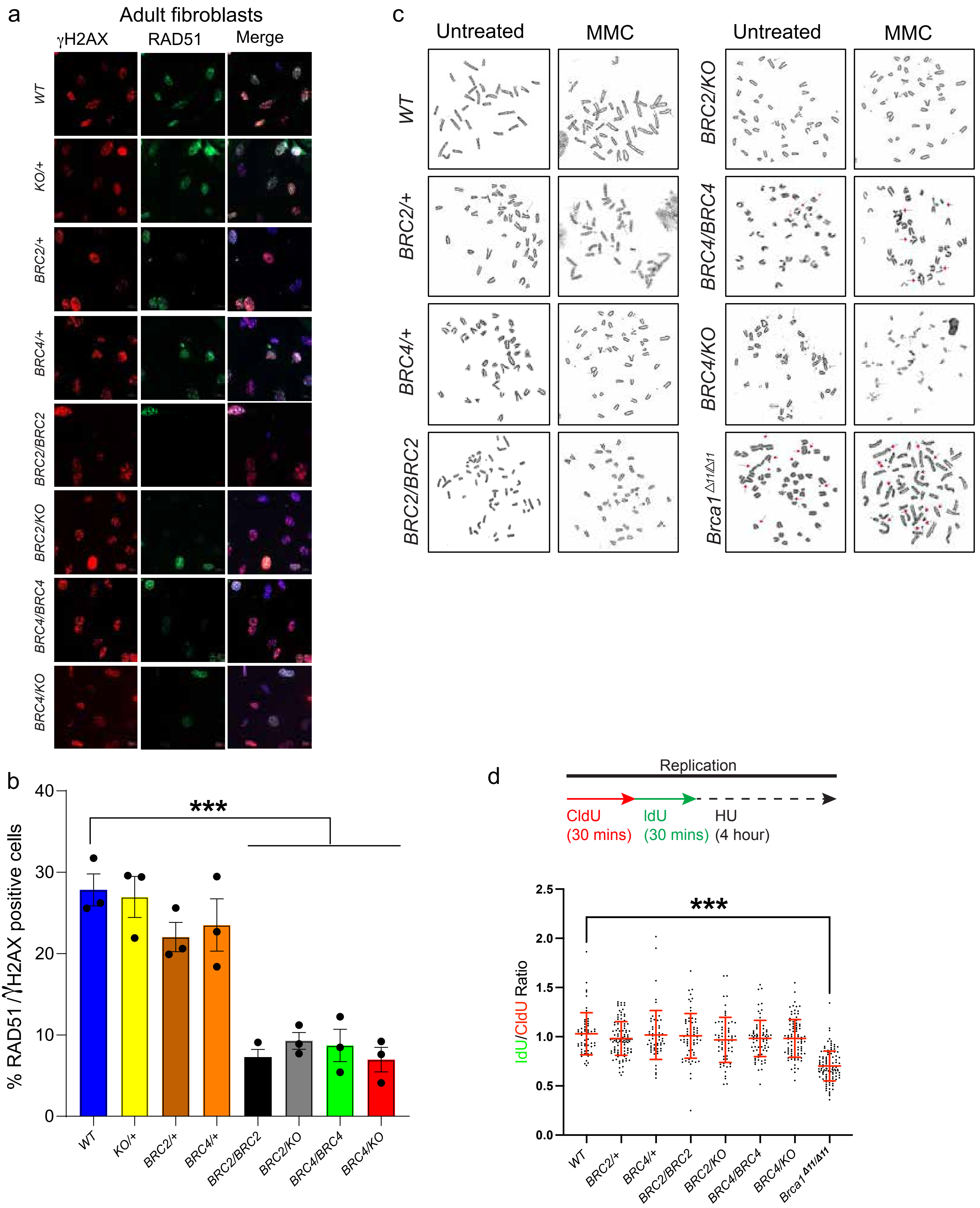

**Supplementary Table 1- List of oligonucleotides used in the study**

| **Name** | **Sequence** | **Size** |
|---|---|---|
| RT-PCR | Forward- agcagatgatgtttcctgtcc<br>Reverse- tttgatttgtgttttcactgtctg | 398bp (for BAC expression) |
| For genotyping BRC mice | FP1- acactctttccctacacgacgctcttccgatct AGGACCAAAAGGCTCACCT<br>FP2- acactctttccctacacgacgctcttccgatct ATTCTCTGGATTTAGCACTGCA<br>RP- gactggagttcagacgtgtgctcttccgatct CCAACTGTGTGTCTTGGTTTCT | 540bp (Mutant)<br>400bp (WT) |
| Genotyping *Mlh1* mutant mice | Mlh1F-tgtcaataggctgccctagg<br>Mlh1R1-tggaaggattggagctacgg<br>Mlh1R2-ttttcagagcagcctatgctc | 500bp (Mutant)<br>350bp (WT) |
| Genotyping *Brca2* KO allele | HPRT fwd-acagcatctaagaagttttgttctgtc<br>Exon11 rev-ctcaacagagtaggttctttgg | No band (WT)<br>450bp (KO allele) |
| gRNAs to generate BRC mice | Brca2-upstream guide (fw): agcccaagtgttgattacaa<br>Brca2-downstream guide (rs): tttgatacgtcttcaggtat | PAM=AGG (For *Brca2* Exon11 targeting) |
| Brc2_ssDNA donor oligo | aatatgaactctgaagaacttttcccagacagtgggaataattttgcttttc aagtaactaataaatgcaataagcctgatttaggaagttcagtggaactc caggaagaagacctcagccacacacaagggcctagtctcaagaactc tcccatggcagtagatgaagatgtagatgatgcgcatgcagcccaagt gttgattacatcttgtttttttaccaaatataaatgaaatggagtttggaggat tttgttctgctcttggcacaaagcttagtgtgtctaatgaggctctgagaaa agctatgaaactgttcagtgacattgaaaatcctgaagacgtatcaaaaat acttcctcaaccttgtgctgaaatcagaaccccagaataccctgtaaactc aaaattgcagaaaacctacaatgataaatccagcttaccaagtaattataa agaaagtggttcttcgggcaatactcaatctattgaagtttctctccaactct ctcagatg | 516 nt |
| Brc4_ssDNA donor oligo | aatatgaactctgaagaacttttcccagacagtgggaataattttgcttttca agtaactaataaatgcaataagcctgatttaggaagttcagtggaactcca ggaagaagacctcagccacacacaagggcctagtctcaagaactctcc catggcagtagatgaagatgtagatgatgcgcatgcagcccaagtgttga ttaagttttcatacagctagcgggaaaaaagtcaaaattatgcaggaatcttt ggacaaagtgaaaaaccttttgatgagacacaacctgaagacgtatcaaa aatacttcctcaaccttgtgctgaaatcagaaccccagaataccctgtaaact caaaattgcagaaaacctacaatgataaatccagcttaccaagtaattataaa gaaagtggttcttcgggcaatactcaatctattgaagtttctctccaactctctc agatg | 492 nt |
